# An integral genomic signature approach for tailored cancer targeted therapy using genome-wide sequencing data

**DOI:** 10.1101/2021.02.17.431663

**Authors:** Xiaosong Wang, Sanghoon Lee, Gong Tang, Yue Wang

**Affiliations:** UPMC Hillman Cancer Center, University of Pittsburgh, Pittsburgh, PA, 15213, U.S.A.; Department of Pathology, University of Pittsburgh, Pittsburgh, PA, 15213, U.S.A.; Department of Biomedical Informatics, University of Pittsburgh, Pittsburgh, PA, 15206, U.S.A.; Department of Biostatistics, University of Pittsburgh, Pittsburgh, PA, 15261, U.S.A.; Department of Surgery, University of Pittsburgh, Pittsburgh, PA, 15213, U.S.A.

## Abstract

With the advent of low-cost sequencing, transcriptome and genome sequencing is expected to become clinical routine and transform precision oncology within next decade. However, viable genome-wide modeling methods that can facilitate rational selection of patients for tailored intervention while tolerating sequencing biases are far lacking. Here we propose an integral genomic signature (iGenSig) analysis as a new class of transparent, interpretable, and resilient methods for precision oncology based on multiple types of genome-wide sequencing data. We postulate that the redundant high-dimensional genomic features, which are typically eliminated during multi-omics modeling, may help overcome the sequencing biases. We thus conceive a novel method that models the therapeutic response using the high-dimensional transcriptional and mutational features predictive of tumor response, which we termed as an integral genomic signature (iGenSig), and then algorithmically resolve the feature redundancy tailored for each patient subject. Using genomic dataset of chemical perturbations, we developed the iGenSig models for predicting targeted therapy responses, and applied selected models to independent datasets for cancer cell lines, patient-derived xenografts, and patient subjects. iGenSig models exhibit outstanding cross-dataset performance compared to artificial intelligence methods, with exceptional resilience against simulated errors in genomic features. In particular, the iGenSig model for the EGFR inhibitor Erlotinib significantly predicted the responses of patient-derived xenografts and patients from a clinical trial, biological interpretation of which led to new insights into the predictive signature pathways with clinical relevance. Together, iGenSig will provide a computational infrastructure to empower tailored cancer intervention based on genome-wide sequencing data.

## BACKGROUND

Precision oncology, defined as molecular profiling of tumors to achieve customized patient care, has entered the mainstream of cancer patient care^1^. The current standard practices for precision oncology include detecting actionable mutations via genetic testing (i.e. EGFR mutation, ALK rearrangements), or detecting small-sized predictive or prognostic gene signatures via targeted expression assays (i.e. Oncotype DX, MammaPrint). Such assays, however, require at least one assay per decision, which limit their cost-effectiveness. On the other hand, the past ten years have observed stunning reduction of sequencing cost for a human genome from $300,000 to $1000, with $100 whole genome sequencing expected soon^2^. With this rate, it is expected that transcriptome and genome sequencing will become the clinical routine for patients. With the advent of low-cost genome sequencing, precision oncology is at the cusp of a deep transformation via leveraging the big data to provide a wide array of clinical decision supports which is deemed to be cost-effective. On the other hand, the computational approaches that can leverage these big data to facilitate clinical decisions and provide tailored health care are far lacking. For example, in metastatic lung cancer, the target therapies prescribed based on the current modeling of genomic sequencing data produced only minimal gain of quality-adjusted life year^3^. Innovative and robust clinical big data-based decision support models for precision oncology will be of vital importance.

In recent years, there has been great enthusiasm about the potential of artificial-intelligence based clinical decision support systems for big data based precision medicine, however, to date only few examples exist that impact clinical practice^4^. The main challenge is that, multi-OMIC big data typically contain daunting amounts of high-dimensional features but limited number of subjects which pose great challenges to the computational power and training process of artificial intelligence (AI) -based methods. In addition, AI approaches are “black box” tools, so that the algorithmic and biological mechanisms underlying the models are largely unknown. The modeling process is controlled by AI which make it difficult to interpret complex model predictions and is often plagued with the problems of overfitting and overweighing. In addition, there is a lack of big-data based methods specifically addressing the insufficient performance of the prediction models for crossing dataset modeling resulting from the common biases in detected genomic features across different datasets arising from sequencing errors, different library preparation methods and platforms, discordant sequencing depth and read-length, heterogenous sample qualities, and experimental variations etc. This calls for robust, transparent, and explainable methods that can predict clinical treatment outcome from multi-OMIC data with substantially improved resilience against sequencing biases.

Here we propose a new class of methods for big data-based precision medicine called integral genomic signature (iGenSig) analysis, which is designed to provide more robust clinical decision support with higher transparency, outstanding resilience and cross-dataset applicability (**Figure 1**). Due to the high dimensionality of genomic features, a common practice for big data-based modeling is to reduce the dimensionality of genomic features via removing redundant variables highly correlative with each other as for gene expression signature panels, or creating synthetic features as for machine learning approaches^5^ (**Supplementary Figure 1**). Here we propose that the redundancies within high-dimensional features can in fact overcome sequencing errors and bias especially when there is a loss of detection of a subset of correlates. Here we define the genomic features significantly predicting a clinical phenotype (such as therapeutic response) as genomic correlates, and an integral genomic signature as the integral set of redundant high-dimensional genomic correlates for a given clinical phenotype such as therapeutic response. The iGenSig analysis generates prediction scores based on the set of redundant genomic features, and then reduce the effect of feature redundancy via adaptively penalizing the redundant features detected in specific samples based on their co-occurrence assessed using large cohorts of human cancers. This allows to preserve the redundant features during the modeling while preventing the feature redundancy from flattening the scoring system. With this method, if a subset of the genomic features was lost due to sequencing biases or experimental variations, the redundant genomic features will help sustain the prediction score. On the other hand, the model utilizes the average correlation intensities of significant genomic features detected in specific samples to diminish the effect of false positive detection resulting from sequencing errors and overweighing. This method also prevents overfitting through dynamically adjusting the feature weights for training subjects. Thus, iGenSig is a simple, white box solution with an integral design to tolerate sequencing errors and bias for big data-based precision medicine. This approach will be more interpretable and controllable than most machine learning or deep learning approaches and will prevent known issues for AI based prediction modeling based on multi-omics big data.

**Figure 1.**
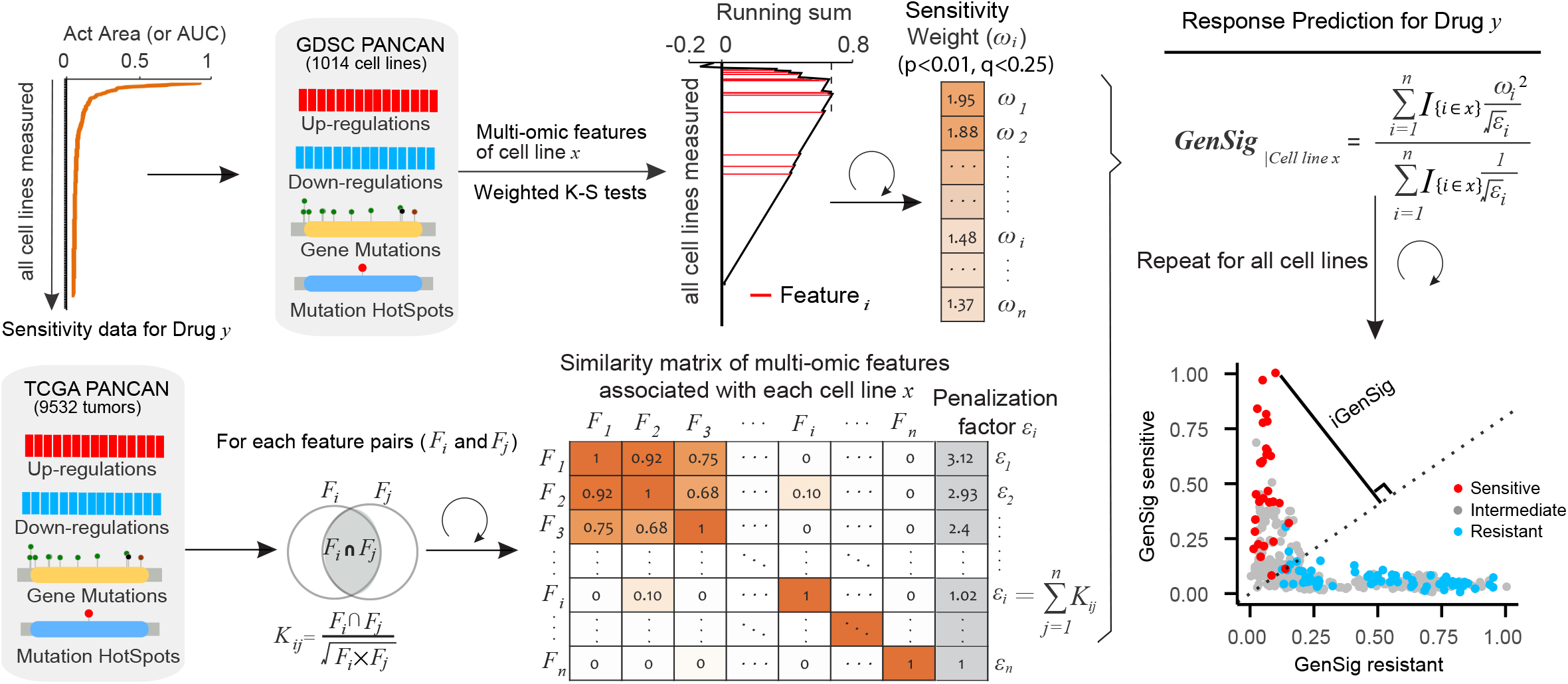
The principle of integral genomic signature analysis. The upper panel shows the calculation of the weights for significant genomic features that predict drug sensitivity or resistance based on weighted K-S tests of Act Area or AUC for each drug, and the lower panel shows the computation of a similarity matrix for genomic features based on TCGA Pan-Cancer dataset to penalize the redundancy between the genomic features associated with each cell line *x*. The resulting sensitive or resistant genomic signature scores are calculated using the indicated formula based on the K-S tests using Act Area or AUC respectively. In the dot plot of sensitive and resistant genomic signature scores for all cell line subjects, a best distinction line (D-line) for separating true sensitive from resistant cell lines is calculated, and the distance of each cell line to this D-line is defined as “iGenSig” score, with positive and negative values indicating sensitive and resistant response predictions respectively.

## RESULTS

To develop the iGenSig modeling, we utilized the drug sensitivity measurements of chemical perturbations, gene expression profiling data, and exome sequencing data for 989 cancer cell lines released by Genomic Datasets of Drug Sensitivity (GDSC, **Supplementary Table 1**). For the drug response measurements, we used high Act Area, the area above the fitted dose response curve (or 1-AUC), to define a sensitive drug response, and high AUC, the area under the dose curve, to define a resistant response. According to literature, the AUC and Act Area are much better quantifiers of drug responses than IC50^6^. To uniform multi-OMIC features, we formulated a Genomic-feature Matrix Transposed (GMT) format for compiling binary multi-OMIC features, similar to that used for compiling gene concepts^7, 8^. Using this format, we analyzed the expression profiling data and exome sequencing data from GDSC and compiled an integrated dataset combining the genomic features including upregulated genes, downregulated genes, mutated genes, and mutation hotspots. We then selected significant genomic correlates using a weighted Kolmogorov–Smirnov (K-S) test that ranks the enrichment of each genomic feature in the cell line panel sorted decreasingly by Act Area or AUC, similar to that implemented by Gene Set Enrichment Analysis (GSEA)^9^. Next, we leveraged the TCGA Pan-Cancer RNAseq and exome dataset for 9532 tumors to quantify the cooccurrence between genomic features associated with each cell line based on similarity measures, which were then used to calculate a redundancy penalty score for each genomic feature.

To prevent the bias from overfitting, we used a random collection of 80% GDSC cell lines as train set and the rest 20% as internal test set for assessing the performance of the model. A total of five train/test sets are generated for modeling through random permutations. We then performed iGenSig modeling for 249 drugs that elicit a negatively skewed drug response distribution in cancer cell lines indicating narrow effect of outstanding responses as observed for most targeted therapies, and have at least 20 sensitive cell line subjects indicating the availability of outstanding responders. To benchmark the performance of the models, we discretized the cell lines into drug sensitive and non-sensitive groups based on a water fall method established in a previous study ^10^, and calculated the Area Under ROC Curve (AUROC) for each drug. As a result, 82 drugs showed an AUROC >0.75 on the testing sets, and 11 drugs showed an ROCAUC>0.85 (**Supplementary Table 2**). Most of the best performing drug models are for targeted therapies against well-known cancer targets such as ERBBs, BRAF, MEK, CDKs, SRC, ATM, TAK, BRDs, PI3K AURORA, BCL2, ABL1, HDAC, AKTs, etc. Among these Lapatinib has the top performing iGenSig model with an average AUROC of 0.95 (**Figure 2a**). The iGenSig scores negatively correlate with the AUC drug measurements in cell lines with a similar trend in both training and testing sets suggesting that iGenSig modeling do not overfit toward training set as opposed to AI-based methods. Among the 249 drugs we modeled, the predictive powers of the iGenSig models appear to significantly correlate with the number of available genomic correlates for each drug (Pearson R=0.59, **Figure 2b, left**), suggesting that the iGenSig models rely on the available genomic information that can predict the drug responses. In addition, the predictive power also significantly correlates with the distribution of cell line responders as indicated by the high GINI scores (Pearson R=0.35, p < 0.01, **Figure 2b, right**). GINI score is used in economy to gauge wealth distribution among a population which is leveraged here to measure the distribution of outstanding responses (**Supplementary Fig. 2a**). In addition, the GINI scores of GDSC drugs do not correlate with the average number of genomic correlates for each drug, suggesting that these are independent factors determining the outcome of iGenSig modeling (**Supplementary Fig. 2b**). Next, we clustered the drugs target kinase signaling based their iGenSig scores in GDSC cell lines, which resulted in more distinctive clustering of the drugs targeting the same or similar kinases (**Figure 2c**) compared to the original drug response measurements (**Supplementary Fig. 3**). Interestingly, outstanding response predictions for BRAF/MEK inhibitors are preferentially enriched in melanoma cell lines, while other drugs such as EGFR inhibitors exhibit cancer type agnostic iGenSig scores, consistent with the tumor-type related clinical activities of these drugs.

**Figure 2.**
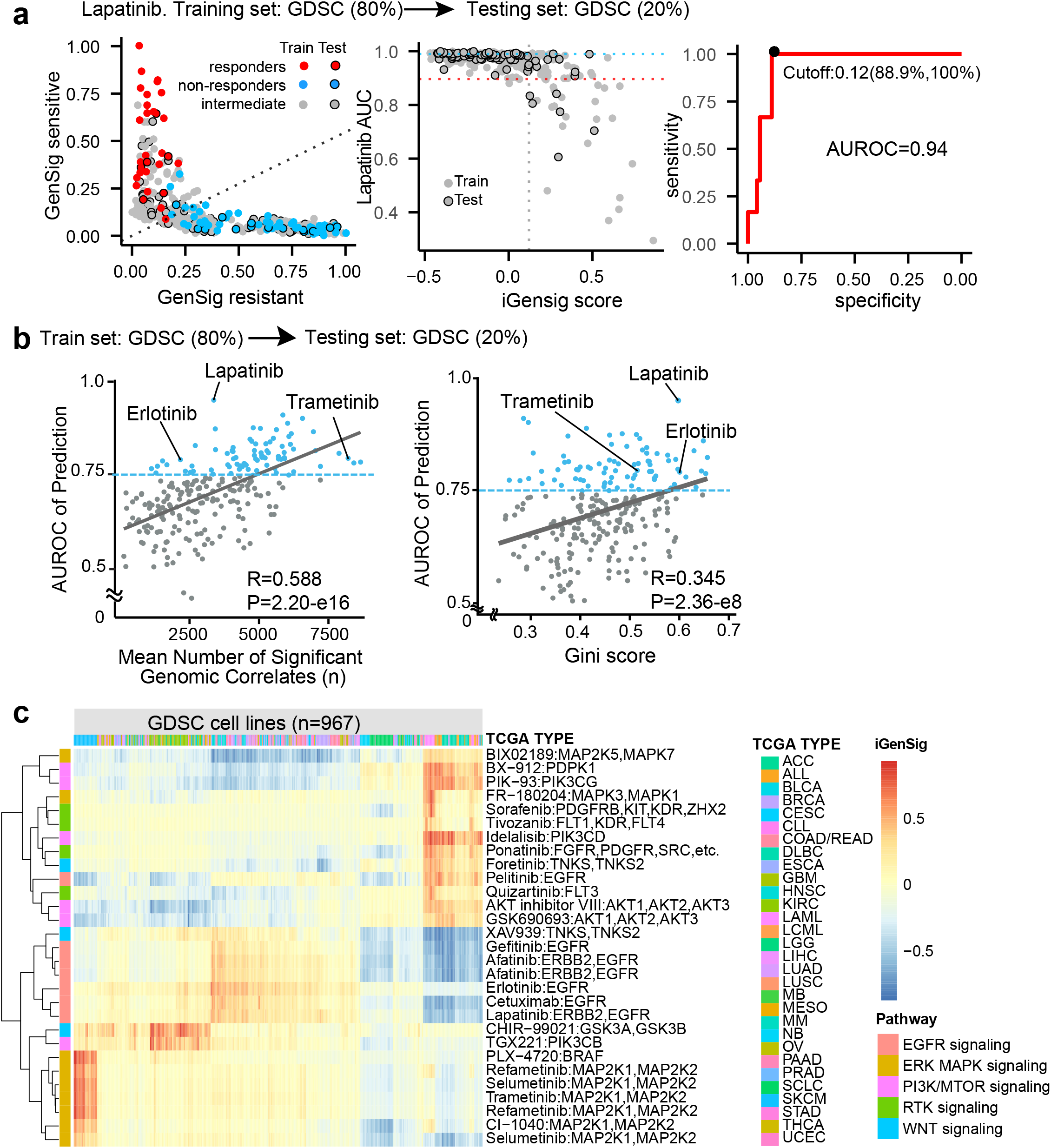
The performance of iGenSig models in predicting the drug responses of GDSC cell lines. **a)** the performance of the iGenSig model for Lapatinib in predicting the response of GDSC cell lines based on a representative training and testing set. Left, sensitive and resistant GenSig scores for GDSC cell lines. Middle, the correlation of the iGenSig scores with AUC measurements for Lapatinib. Right, the receiver operating characteristic (ROC) curve for predicting sensitive responses to Lapatinib. As golden standard for the ROC curve, the cell line subjects in the test set are divided into sensitive and non-sensitive groups based on the AUC measurements for Lapatinib using the cutoff determined by the waterfall method (see Methods). **b**) the performance of the iGenSig models for 249 drugs assessed by their average AUROC in correlation with their average number of significant genomic features (left) or their GINI scores (right). The average AUROC for each drug was calculated based on five train/test sets. **c**) clustering GDSC cancer cell lines and targeted kinase drugs based on iGenSig scores. The drugs targeting different kinases or different kinase families form distinctive clusters.

To assess the cross-dataset performance of our iGenSig models, we analyzed the RNAseq and exome sequencing data from the Cancer Cell Line Encyclopedia (CCLE). In total there are eleven drugs measured by both CCLE and GDSC datasets. The drug response measurements by GDSC and CCLE modestly correlate with each other (**Figure 3a**), consistent with the previous report^11^. Our result showed that the predictive performance of iGenSig models on the CCLE dataset appear to correlate with their performance on the testing sets of the GDSC dataset (Pearson R=0.78, **Figure 3b**). More interestingly, the iGenSig scores calculated based on GDSC models showed more obvious correlations with the Act Areas of CCLE cell lines than the GDSC AUC measurements (**Figure 3a, c**). Using GDSC as training set and CCLE as validation set, the models for these three drugs, Lapatinib, Erlotinib, and Sorafenib, achieved a prediction AUROC of 0.85, 0.81, and 0.80 respectively. Plotting the significant genomic features for Erlotinib in the two datasets revealed consistent integral genomic signature correlating with drug sensitive or resistant responses (**Figure 3d**). This suggest that the modest consistency between the GDSC and CCLE measurements could be attributed to the number of cell lines screened by both GDSC and CCLE for which insufficient sensitive cell lines were screened in both projects, as suggested by the previously study^11^, or due to the different cellular states under different cell culture conditions.

**Figure 3.**
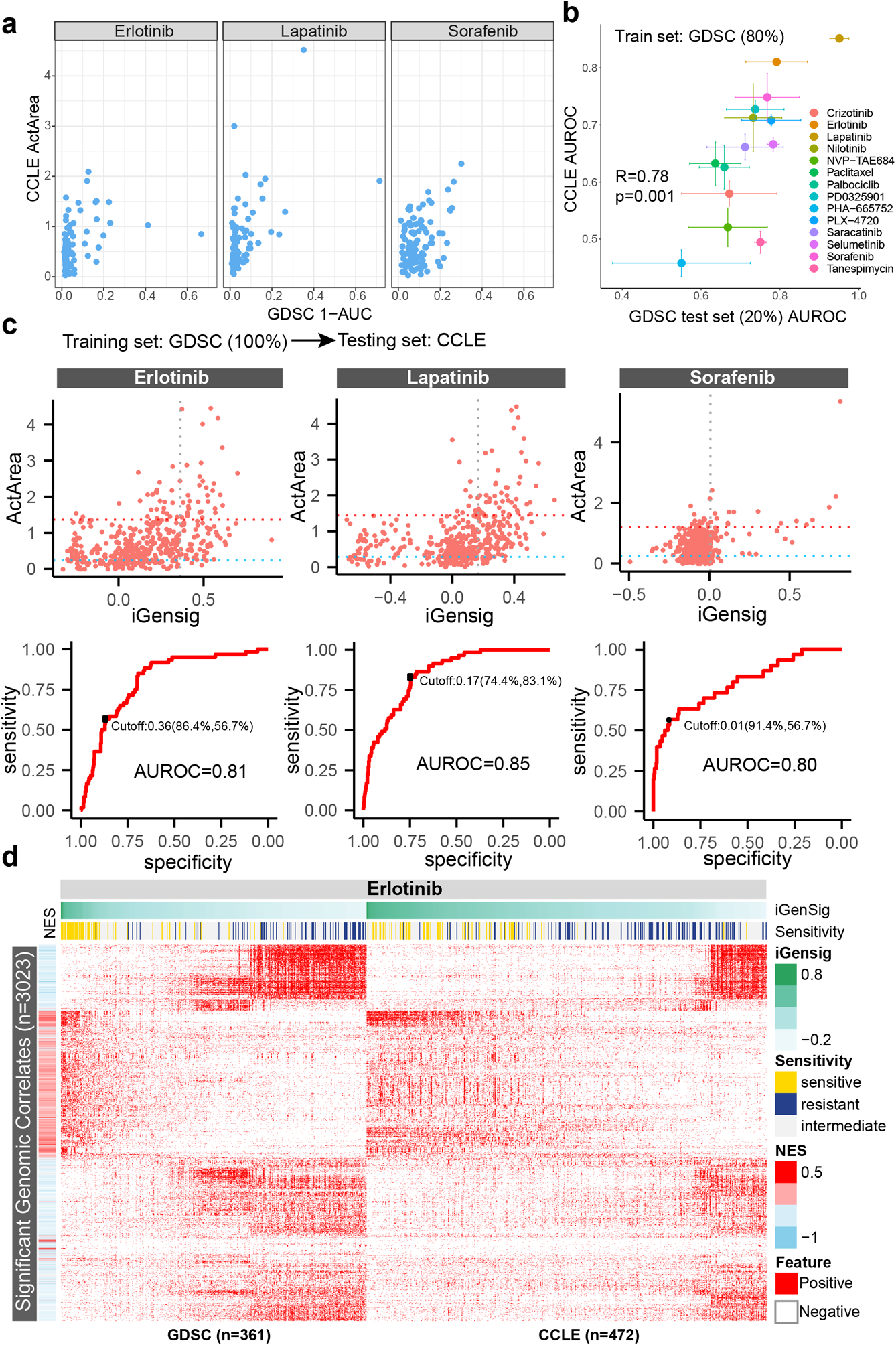
Predictive values of iGenSig models developed from GDSC phamacogenomic dataset on the CCLE cell line responses to the respective drugs. **a)** the correlation between the act areas of cell lines measured by both GDSC and CCLE for Erlotinib, Lapatinib, and sorafenib. **b)** the performance of GDSC iGenSig models in predicting the drug responses of GDSC testing cell lines and CCLE cell lines. 80% of GDSC cell lines are used for buiding the iGenSig models and 20% of GDSC cell lines are used for testing. 100% of CCLE cell lines are used for cross-dataset validations. **c**) The predictive values of the iGenSig models developed from GDSC data on the CCLE data for Erlotinib, Lapatinib, and Sorafenib. Upper panel shows the correlation between the iGenSig scores and the Act Areas of the respective drugs for CCLE cell lines. The horizontal dashed lines shows the cut offs for sensitive (red) and resistant (blue) calls. The vertial dashed line shows the optimal cut off for iGenSig scores determined based on AUROC. The lower panel shows the ROC curves of iGenSig scores in determining the sensitive cell lines vs non-sensitive cell lines. **d)** GDSC and CCLE cell lines show consistent integral genomic signature that correlates with Erlotinib responses. The significant genomic features (n=3023) based on K-S tests are shown in the figure. The GDSC and CCLE cell lines are sorted by their iGenSig scores. The cell lines that have been tested for Erlotinib chemical perturbations are shown in the figure, and the sensitive and resistant cell line subjects are indicated as yellow and blue bars.

To test if the iGenSig predictions rely on the genomic features of the primary drug targets, we removed the drug target genes for Erlotinib, Lapatinib, or Sorafenib from GDSC and CCLE genomic feature sets. We then built the iGenSig models for these drugs based on the genomic features devoid of drug targets and assessed their performance on GDSC internal test set or the CCLE validation set (**Figure 4a**). Our result showed that the performance of the iGenSig models are not affected by the absence of genomic features for known drug targets (**Figure. 4b**). Next, we sought to compare the performance of iGenSig modeling with AI-based methods. Following the previously reports^12, 13^, for dimensionality reduction we computed the unsupervised representation of the genomic features based on the autoencoder deep learning method which were then fed to the machine learning methods for supervised learning on drug responses, such as elastic net, neural network, support vector machine (SVM) or Random Forest (RF) (**Supplementary Fig. 4**). We then applied these methods to model cancer cell sensitivity to Erlotinib, Lapatinib, and Sorafenib. Compared to our iGenSig model, the AI based methods can achieve favorable prediction on GDSC internal test set with average AUROC between 0.89-0.90 except neural network (**Figure 4c**). Among the AI-based methods, the random forest and neural net showed obvious overfitting when its performance is tested on the GDSC training set. However, when applied to CCLE dataset, the AUROCs of prediction for these methods dropped to 0.65-0.82, whereas the iGenSig models maintained significantly higher predictive values (**Figure 4c**). To assess the resilience of these models against the common biases resulting from experimental variations and sequencing errors, we simulated the biases in genomic features by generating false-positive or false-negative genomic features in either GDSC or CCLE dataset. By randomly deleting or inserting genomic features, we generated simulated genomic feature sets with 5-25% error rates (five random samplings were performed for each error rate level) and tested the effects on the iGenSig models and the best-performing AI-models generated using autoencoder and elastic net. Our result showed that the simulated errors in genomic features jeopardized the predictive values of the AI-based models for Lapatinib and Erlotinib, which are particularly vulnerable to simulated errors in the training set (**Figure 4d**). Whereas the respective iGenSig models can tolerate simulated errors for up to 20% of genomic features without obviously decreasing its performance, irrespective of whether the errors are simulated in training or validation sets.

**Figure 4.**
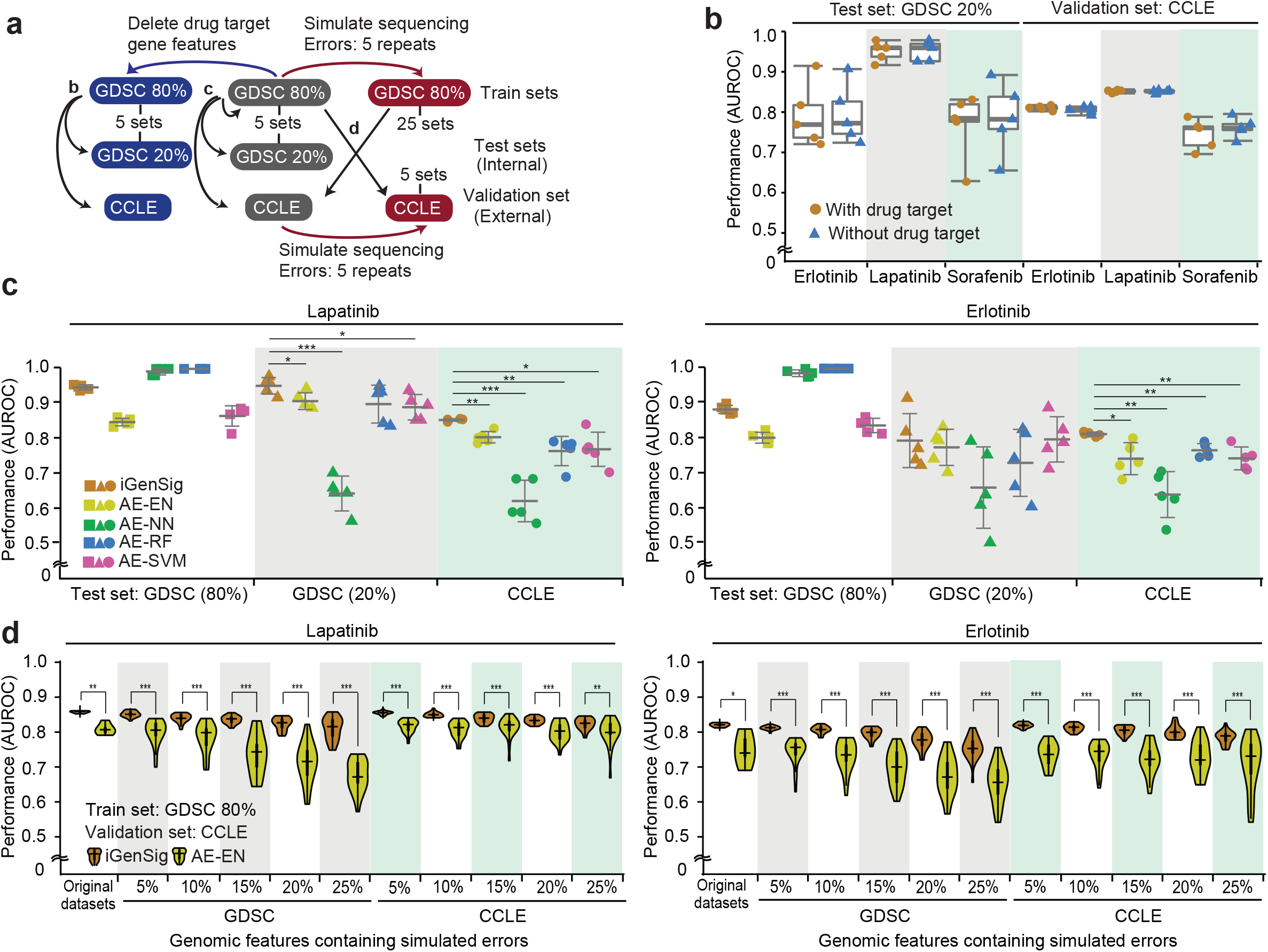
The performance of iGenSig modeling based on genomic features devoid of drug targets or containing simulated sequencing biases, and comparisons between iGenSig modeling and AI-based modeling. **a**) Schematic showing the training, testing and validation sets used for drug sensitivity modeling shown in b-d. **b**) The prediction performance of iGenSig models do not depend on the genomic features derived from the drug target genes. The box plot shows the performance of the iGenSig models for Erlotinib, Lapatinib, and Sorafenib assessed on GDSC testing set (left) or CCLE validation set (right) in the presence or absence of the genomic features derived from the respective drug targets. **c**) Prediction performance of the iGenSig model and AI models for Lapatinib and Erlotinib on the GDSC training and testing sets or the CCLE validation set. For AI methods, the unsupervised learning was performed by autoencoder (AE) and supervised learning was performed using various machine learning tools including elastic net (EN), neural network (NN), random forest (RF) and support vector machine (SVM). The prediction performances were assessed on GDSC 80%, GDSC 20%, or CCLE cell lines. **d**) Comparing the prediction performance of the iGenSig model and the AE-EN machine learning model for Lapatinib and Erlotinib using genomic features containing the specified percentages of simulated errors. Sequencing biases were simulated on GDSC genomic features or CCLE genomic features (left or right side respectively for each drug). The unsupervised learning was performed by autoencoder (AE) and supervised learning was performed using elastic net (EN). * P<0.05, ** P<0.01, and *** P<0.001 (unpaired two-tail t-test).

Next, we sought to test the applicability of the GDSC iGenSig models to predict the therapeutic sensitivity of the patient derived xenograft (PDX) tumors, which are transplanted from human tumors without any *in vitro* manipulation, thus are ideal for testing therapeutic effects. This can be achieved using a published drug sensitivity dataset of PDX tumors with gene expression and mutation data^14^. This dataset measured the sensitivity of PDX tumors against 36 single drugs, among which four drugs are also profiled by GDSC with favorable iGenSig models (AUROC >0.75). Applying our GDSC iGenSig models for these drugs revealed significant predictive values for two drugs on the survival of xenopatients: Erlotinib and Trametinib (**Figure 5a**). Next, we sought to test the performance of our GDSC iGenSig model for predicting patient response to Erlotinib monotherapy. Most clinical trials for these targeted drugs assessed their combinations with chemotherapies instead of monotherapies, which will confound the outcome of drug response modeling. Literature investigation revealed that the dataset of Biomarker-integrated Approaches of Targeted Therapy for Lung Cancer Elimination (BATTLE) trial (GSE33072) profiled non-small cell lung cancer (NSCLC) tumors from 131 patients by gene expression array, among which 28 patients are treated with Erlotinib monotherapy, and 47 patients are treated with Sorafenib monotherapy. Progression free survival analysis suggested that, the GDSC iGenSig model significantly predicted the favorable response of these patients to Erlotinib with a hazard ratio of 0.3 (p=0.026, **Figure 5b**). These results support the utility of integral genomic signature modeling in predicting therapeutic responses of targeted therapies and its outstanding cross-dataset performance.

**Figure 5.**
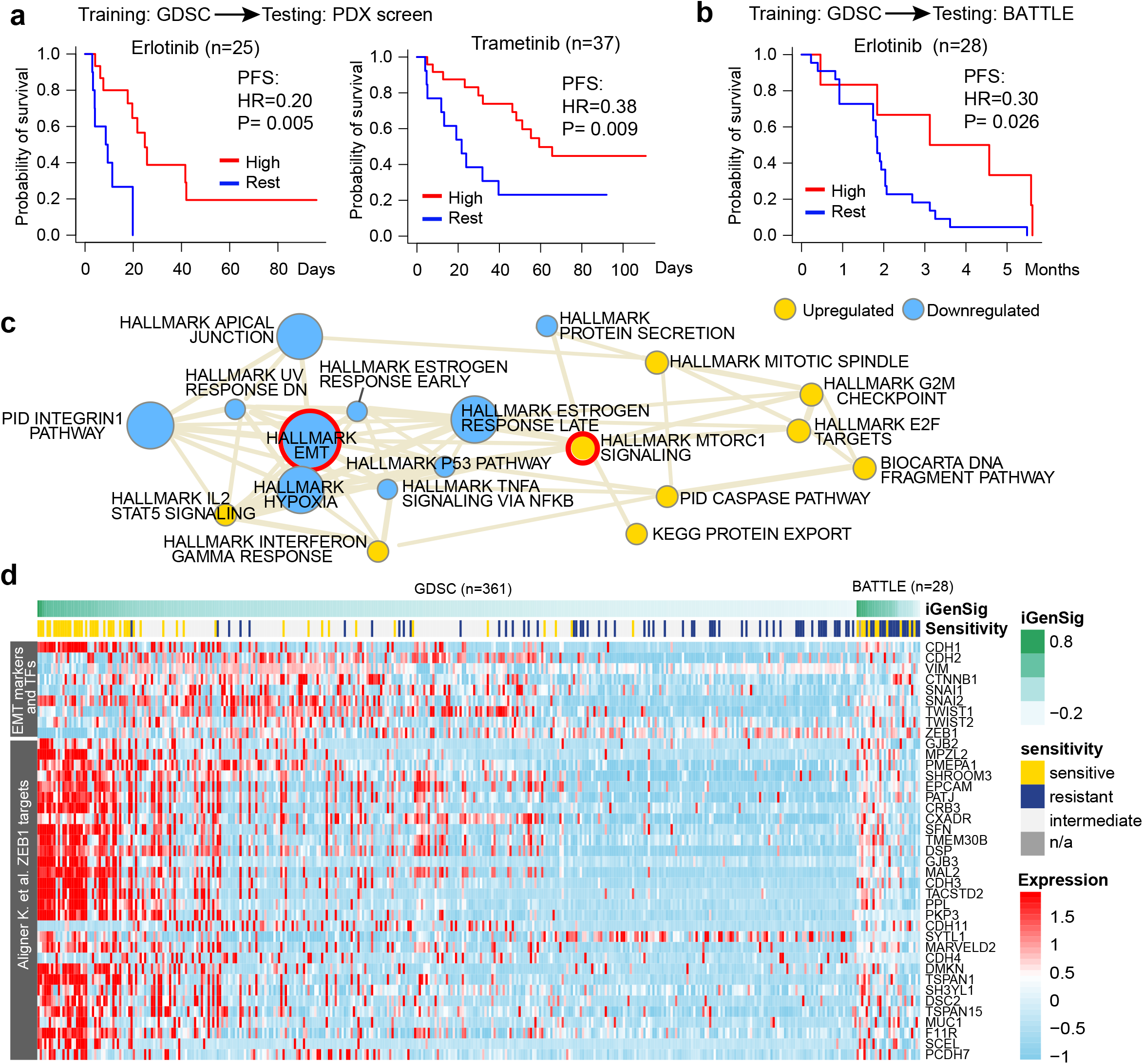
Predictive values of iGenSig models developed from GDSC cell line phamacogenomic data on the survival of xeno-patients from the PDX preclincial trials and human subjects from the BATTLE trial. **a**) Kaplan–Meier plots showing the predictive values of the iGenSig models developed from GDSC dataset on the PDX xenopatient trials for Erlotinib and Trametinib. **b**) Kaplan–Meier plot showing the predictive values of GDSC iGenSig models for Erlotinib on the patients from the BATTLE trial. A data-driven cut point of high iGenSig scores was determined as previously described^29^. P-values are based on Gehan-Breslow-Wilcoxon test (a) or log-rank test (b). (**c**) The network of upregulated and downregulated pathways characteristic of Erolotinib resistant GDSC cell lines and human subjects. The top ten upregualted pathways (gold) and downregulated pathways (blue) that were clustered in the interconnected network are shown in the figure. The CSEA enrichment score for each pathway in the Erlotinib resistant signature is depicted by the size of each node. The pathway associations are depicted by the thickness of the edge. The pathway associations are calculated based on CSEA association scores between each pair of pathway, and the edges with association scores more than 2 are shown in the figure. (**d**) Heatmap showing the associations of EMT markers and master transcription factors as well as ZEB1 target genes with the Erlotinib iGenSig scores in the GDSC and BATTLE datasets. The cell lines and patient subjects are sorted based on their iGenSig scores.

Since epithelial mesenchymal transition (EMT) has been previously reported to mediate EGFR resistance in the BATTLE trial study^15^, we wonder if the EMT signature contribute to the iGenSig predictions. We thus examined the pathways characteristic of the integral genomic signature for Erlotinib resistance in our iGenSig model. This can be achieved by extracting the genes contributing to the resistance related genomic features in our GDSC iGenSig model. The resulting resistance gene list can be then used to explore the enriched pathways based on the concept signature enrichment analysis (CSEA) developed in our previous study, which is designed for deep functional assessment of the pathways enriched in an experimental gene list^8^. Our result showed that the most significantly upregulated pathways characteristic of Erlotinib resistance signature include MTORC1 signaling and E2F target gene signature (**Figure 5c, Supplementary Fig. 5**). Consistent with this, activation of mTOR signaling including activating mutation or overexpression of mTOR has been found to mediate resistance to EGFR inhibitors and predict worse patient outcome in lung cancer^16, 17^. On the other hand, the EMT pathway is ranked as the most significantly downregulated pathways in the Erlotinib resistance signature identified from GDSC cell lines, which contradict with the previous report^15^.

We postulated that this may be attributed to the content of the EMT signature that mixed both upregulated and downregulated genes in EMT. We thus compiled a upregulated EMT signature and a downregulated EMT signature based on a previous report^18^. Correlating these EMT signatures with the Erlotinib iGenSig scores revealed that the downregulated and upregulated EMT signatures are indeed enriched in the subjects with high or low iGenSig scores respectively in the BATTLE trial dataset (**Supplementary Fig. 6-7**). However, in the GDSC Pan-Cancer cell line panel, both upregulated and downregulated EMT signatures are repressed in Erlotinib-resistant cell lines. This suggests that repression of both EMT signatures are characteristic of the Erlotinib resistant cell lines at Pan-cancer scale and explains the pathway results from CSEA analysis. Among the known EMT markers and transcription factors, overexpression of E-cadherin (CDH1) was observed in both sensitive cell lines and patient responders from BATTLE trial (**Figure 5d**). Whereas overexpression of EMT markers such as N-cadherin (CDH2), Vimentin (VIM), and β-catenin (CTNNB1) are characteristic of the cell lines with intermediate sensitivity, and overexpression of ZEB1 is characteristic of Erlotinib resistant cell lines. In the BATTLE trial, overexpression of either β-catenin or ZEB1 are characteristic of subjects with low iGenSig scores. As ZEB1 is a transcriptional repressor, we assessed the correlation of ZEB1 target genes with the iGenSig scores. This revealed that downregulation of ZEB1 target genes are characteristic of both resistant cell lines and patient subjects (**Figure 5d**). Together, our results suggest that EMT is associated with reduced but intermediate response to Erlotinib whereas repression of ZEB1 signature is associated with tumor-type agnostic resistance at Pan-cancer scale.

## Discussion

Here we introduce a new class of integral genomic signature methods that leverage the high-dimensional redundant genomic features as an integral genomic signature to enhance the resilience of multi-omics-based modeling for precision modeling. The iGenSig method is designed to address the transparency, resilience, and interpretability issues for big-data based modeling. In particular, our iGenSig models demonstrated outstanding performances in crossing datasets and tolerating the bias in the detected genomic features. IGenSig models can be managed in every detailed step, and the underlying pathways can be biologically interpreted through the concept signature enrichment analysis we developed^8^. The performance of iGenSig models appears to at least in part depend on the availability of outstanding responders and significant genomic correlates. We expect that iGenSig as a new class of big-data based modeling methods will have broad application in modeling therapeutic responses based on pharmacogenomic and clinical trial datasets. Further, iGenSig can also be potentially applied to predict other cancer behaviors to facilitate clinical decision such as aggressiveness of carcinoma in situ, or metastatic potential of clinically localized tumors.

It is interesting to note that our iGenSig model for Erlotinib demonstrated predictive value in both patient-derived xenografts and the patients in the BATTLE trial. Interpretation of this model revealed that downregulation of ZEB1 target genes are characteristic of Erlotinib resistance signature, whereas induction of EMT is associated with reduced but intermediate responses. While both EMT and ZEB1 has been found to medicate acquired resistance to EGFR inhibitors in non-small cell lung cancer^19, 20^, our iGenSig signature suggested the discordance of ZEB1 overexpression with EMT induction, and their differential contribution to Erlotinib resistance in Pan-cancer cell lines. This implies that the phenotypic changes other than EMT induced by ZEB1 may contribute to Erlotinib resistance. Consistent with this finding, ZEB1 has been reported to exert more critical functional consequence than EMT itself. For example it has been proposed that specific EMT inducers such as ZEB1, but not the EMT state, that determine cancer stem cell properties^21^. Future experimental studies will be required to determine the differential contribution of ZEB1 and EMT to Erlotinib resistance in human cancers.

On the other hand, our GDSC iGenSig model for sorafenib failed to predict patient response in the sorafenib treatment arm in the BATTLE trial dataset. To explore if this is attributable to the differences in the *in vitro* and *in vivo* microenvironments, we examined the primary targets of this drug. Sorafenib is a multi-kinase inhibitor of Raf-1, B-Raf and VEGFR-2. Among these, VEGFR-2 is an VEGF receptor involved in angiogenesis. In light of this, we wonder if the iGenSig models based on *in vitro* cell line responses cannot model *in vivo* tumor responses to VEGFR inhibition. Based on literature, VEGFA amplification is a known biomarker for Sorafenib response^22^, whereas CXCL8 (IL8) are known to induces VEGF overexpression in endothelial cells and promote angiogenesis^23^. Correlating these biomarkers with the iGenSig scores revealed that in the BATTLE trial dataset, the sensitive tumors with low iGenSig scores appear to overexpress VEGFA and CXCL8 (**Supplementary Fig. 8**). This suggests that the inability of GDSC iGenSig model to predict patient response to sorafenib in the BATTLE trial may be attributed to the anti-angiogenesis activity of sorafenib, which cannot be modeled using *in vitro* cultured cell lines. This reflects the limitation of modeling patient tumor response based on *in vitro* cell line models.

The remaining issue to be addressed for iGenSig modeling is how to eliminate the effect of confounding genomic features resulting from imbalanced distribution of confounding factors such as gender or prognostic factors that can impact patient outcome such metastasis, etc. While this issue may be less impactful in modeling of cell line responses when a large number of cell line subjects are included, it could become more consequential when a smaller number of subjects are tested in the clinical trial. In this case, the genomic features associated with the confounding factor may be identified and excluded from the iGenSig model through multi-variate statistics. In addition, the confounding clinical variables that affect prognosis such as metastasis should be accounted for via multivariate statistics during the iGenSig modeling based on survival outcome. Future studies will be required to further optimize the iGenSig methods for modeling clinical trial datasets and taking into consideration of these biological variables and confounding factors.

## Materials and Methods

### Data retrieval

The drug response data, gene expression data, and mutation data are from the Genomics of Drug Sensitivity in Cancer Project (GDSC), and the Cancer Cell Line Encyclopedia (CCLE) as of September 2018. The GDSC and CCLE gene expression data are retrieved from ArrayExpress (E-MTAB-783) and NCBI GEO (GSE36133) respectively, and normalized using Robust Multi-Array Averaging (RMA)^24^. Drug sensitivity data, mutation data, and cell line annotations were downloaded from the GDSC (http://www.cancerrxgene.org/downloads) or CCLE (http://www.broadinstitute.org/ccle) websites. The gene expression and mutation data for the PDX tumors included in high-throughput drug response screening were retrieved from the supplementary dataset of the original publication^14^. The TCGA Pan-cancer gene expression and mutation datasets were retrieved from UCSC Xena browser (https://xenabrowser.net). The gene expression data for Biomarker-integrated Approaches of Targeted Therapy for Lung Cancer Elimination (BATTLE) trial were retrieved from GEO (GSE33072).

### Extracting genome-wide expression and mutation features for cell line and tumors

Based on gene expression and somatic point mutation datasets, we extracted genome-wide gene expression and mutation features and generated an integrated genomic feature file. For gene expression data, we first calculated log2 transformed fold changes of the expression values compared to the trimmed mean of expression values (excluding the 10% largest and 10% smallest values). To eliminate zero values during log2 transformation, we assigned the zero values as half of the minimum expression value across all cell lines or tumors. Based on the mean and standard deviation (SD) of fold changes, we assigned the cell lines or tumors into the following overlapping groups: ‘Up_Level1’ group with the fold change above Mean+1*SD for a given gene; ‘Up_Level2’ group with the fold change above Mean+2*SD; and ‘Up_Level3’ group with the fold change above Mean+3*SD Likewise, ‘Down_Level1’, ‘Down_Level2’, and ‘Down_Level3’ grouped cell lines based on Mean-1*SD, Mean-2*SD, and Mean-3*SD. The 6 ‘Levels’ were labeled as genotypic features for each given gene and the binary genomic features are compiled as a Genomic Matrix Transposed (GMT) file format. Similarly, we extracted binary genomic features to represent point mutations. The mutation hotspots and nonsynonymous somatic mutations such as missense, nonsense, and frame shift are assigned as mutation features. Each recurrent mutation hotspot and each recurrently mutated gene were assigned as separate features.

### Defining drug responses of cancer cell lines

Drug responses of cancer cell lines are represented by the area under dose-response curve (AUC) in GDSC or the area over the dose-response curve (Act Area) in CCLE^10, 25^. We first tested the skewness of the AUC measurements for each drug in the GDSC dataset. A negative skewness distribution indicates that the drug has high AUC measurements (lack of responses) in most of the cell lines, but low AUC measurements (sensitive responses) in a small subset of the cell lines, and a lower level of skewness indicates higher level of outstanding responses. To ensure the drugs have sufficient outstanding responders for training and testing the algorithm, the GDSC drugs with negative skewness and more than 20 sensitive cell line subjects are included in our iGenSig modeling. We then defined sensitive drug responses of cell lines based on Act Areas using the water fall method described in the CCLE study^10^. The Act Area measurements for CCLE or GDSC cell lines for a given compound are sorted in ascending order to generate a waterfall distribution. The cut-off for defining sensitive subjects was determined as the maximal distance to a line drawn between the start and endpoints of the distribution. The cut-off for non-responders was determined as ‘median of Act Area - median absolute deviation (MAD).’ The cell lines with Act Area above the sensitivity cut-off were labeled as drug-sensitive and below the resistance cut-off were labeled as drug-resistant. The cell lines with Act Areas between the cut-offs for drug sensitivity and resistance were labeled as intermediate.

### The basic algorithm for calculating genomic signature scores

To define the weight (ω_*i*_) of each genomic feature in predicting sensitive drug responses, we leveraged the weighted Kolmogorov–Smirnov (WKS) statistics^9^ to test the enrichment of the feature-positive cell line in the cell line panel sorted in descending order based on Act Area (**Figure 1**). To prevent bias, we excluded the genomic features defining fewer than 5 cell lines during the calculation of GenSig scores. Likewise, we calculated the weights for each genomic feature in predicting resistant drug responses based on the cell line panel sorted by AUC in descending order. We assessed the significance of the observed enrichment score (*ES*) by comparing that to the random ES scores calculated by random features with the same numbers of positive cell lines. Repeat this step until 1000 random enrichment scores were calculated, then the normalized enrichment score (NES) was calculated by:

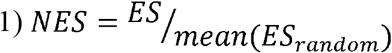

The p-values were determined based on the chance of random ES scores to be above the observed ES score for feature *i*, and the false discovery rate (FDR) were calculated using R package “qvalue”. Significant genomic features with a p-value<0.01 and FDR q-value<0.25 were selected for calculating GenSig scores.

To prevent the inflation of iGenSig scores from feature redundancy, we leveraged the TCGA Pan-Cancer RNA-seq and exome datasets to assess the co-occurrence between genomic features associated with each cell line and generated the cosine similarity matrix of genomic features based on Otsuka-Ochiai coefficient between these features (K_ij_). We then introduced a penalization factor (ε) calculated based on the similarity matrix of genomic features associated with a given cell line _*x*_.

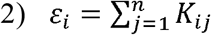

Where n is the total number of genotypes associated with cell line _*x*,_ K_ij is_ the Otsuka-Ochiai coefficient between each pair of the genotypes associated with cell line _*x*._ To eliminate the cumulative effect of small overlaps between genomic features, the Otsuka-Ochiai coefficients were adjusted to 0 if K_ij_<0.1. Here ε_*i*_ is an estimator of redundancy among the genomic features associated with cell line _*x*._ The penalization factor ranges from 1 (all genotypes are completely different each other) to n (all genotypes are the same). We then penalized the weight ω_*i*_ using the square root of ε_*i*_, resulting in Effective Weight (EW):

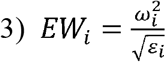

The sum of the reciprocals of square root ε_*i*_ was then used to calculate the Effective Feature Number (EFN):

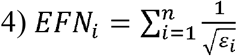

Finally, the GenSig score of the given cell line_x_ is computed as:

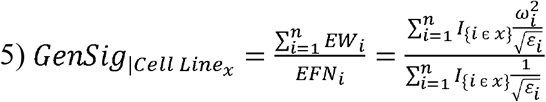

### Calculating iGenSig scores for predicting drug sensitivity and resistance

For a given drug, based on WKS tests using either Act Area or AUC, we calculated sensitive or resistant GenSig scores for each GDSC cell line. We then sorted all GDSC cell lines based on the ratio of sensitive and resistant GenSig scores for a given drug decreasingly. Scanning each cell line from the top to the bottom of this sorted cell line list, the percentage of sensitive cell lines ranked higher than each cell line is calculated as:

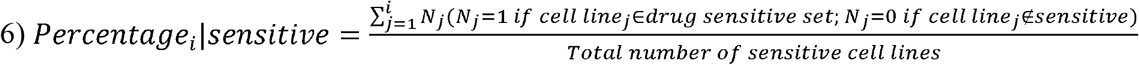

Similarly, the percentage of resistant cell lines ranked higher than each cell line is calculated as:

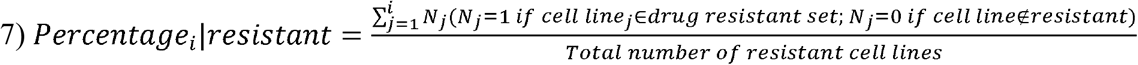

Then the Youden Index for determining the optimal ratio to separate sensitive from resistant cell lines was calculated as below:

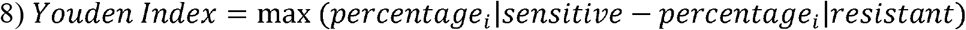

In a two-dimensional plot by the *GenSig*|*sensitive* and *GenSig*|*resistant* scores, cell lines will be placed according to the *GenSig*|*sensitive* and *GenSig*|*resistant* scores. The slope of the dividing line (D-line) for sensitive and resistant cell lines are determined based on Youden Index. With this computation, if a cell line has high probability to be drug sensitive, it will have higher *GenSig*|*sensitive* score than *GenSig*|*resistant* score and will be placed above the D-line. Therefore, the possibility for a cell line to be sensitive or resistant can be assessed by their vicinity to the D-line (**Figure 1**). We thus calculated the distance between a cell line and D-line and defined the distance as the final iGenSig score, which can be used to predict its drug sensitivity. The sensitive cell lines above D-line will have positive iGenSig scores and vice versa.

### Benchmarking the performance of the iGenSig algorithm

To benchmark the performance of the iGenSig algorithm in determining drug sensitivity, we randomly selected 20 % of GDSC cell lines treated by a specific drug as internal test set. We assigned the rest of 80% cell lines as train set and performed this randomized sampling for 5 times. The distributions of drug sensitive and resistant cell lines were required to be balanced between the train and test set in each sampling. The CCLE dataset was used as external validation set of our predictive models to assess their applicability to an independent dataset. The area under ROC curve (AUROC) of the iGenSig scores was calculated based on the binary response of the cell lines determined based on the sensitive cutoff discussed above. The cell line subjects were divided into sensitive cell lines and other cell lines that include both intermediate and resistant cell lines.

To test if the iGenSig predictions rely on the genomic features of the primary drug targets, we removed the drug target gene features for Erlotinib, Lapatinib, or Sorafenib from GDSC and CCLE genomic feature sets. We then built the iGenSig models based on the genomic features devoid of drug targets and assessed their performance on internal test set (GDSC 20% cell lines) or external validation set (CCLE cell lines). To simulate the common biases in genomic features resulting from experimental variations or sequencing errors, we generated false-positive or false-negative genomic features in either GDSC or CCLE dataset. By randomly deleting or inserting genomic features, we generated simulated genomic feature sets with 5, 10, 15, 20, or 25% error rates and repeated the simulations for five times for each error rate, which were applied to the five randomized GDSC training sets or the CCLE validation set (**Figure 4a**). We then applied the same sets of genomic features with or without error simulations to iGenSig modeling, or the deep learning and machine learning modeling methods discussed below to compare the resilience of iGenSig models with the AI-based methods.

### Deep learning and machine learning methods

Following the previously reports^12, 13^, we applied the deep learning method autoencoder^26^ to perform unsupervised representation learning for dimensionality reduction and machine learning prediction algorithms for supervised learning of therapeutic responses using the low dimensional features generated by autoencoder and compared their prediction performances with our iGenSig method. The Autoencoder model was developed using the same genome-wide gene expression and mutation features we compiled, and we used the same training, internal testing, and external validation sets of cell line models as in iGenSig modeling. The autoencoder model was built with three hidden layers and the unit sizes 150, 50, and 25 in each layer with the “ReLU” activation function and encoded synthetic feature size of 10. We then applied the unsupervised representation of the genomic correlates to supervised learning methods including elastic net, artificial neural network, Random Forest (RF), and support vector machine (SVM) for prediction modeling. Elastic net is a regression method that combines lasso and ridge regularization with the two hyperparameters, alpha and lambda. Alpha is a mixing parameter to define the relative weight of the lasso and ridge penalization terms and lambda determines the amount of shrinkage^27^. We identified alpha with the best tuning and optimized for predictive performance over a range of lambdas. Regression was performed using the glmnet R package (ver. 4.0.2). Artificial neural network is non-linear learning model and contains hidden layers that use backpropagation to optimize the weights of the input variables to improve the predictive power of the model. For artificial neural network prediction, we specified three hidden layers with 5, 3, and 10 nodes and 0.01 threshold for the partial derivatives of the error function using the neuralnet R package (ver. 1.44.2). We implemented RF regression model using randomForest R package (ver.4.6.14). We specified 1,000 trees to grow and ensure every object gets predicted multiple times. We used SVM with linear kernel method, ‘svmLinear2’, provided by caret R package (ver. 6.0.86). We specified tuneLength=10 in the tuning parameter grid and accuracy metric.

### Pathway enrichment analysis for integral genomic signature

To identify the pathways characteristic of the integral genomic signature for Erlotinib resistance modeled from the GDSC dataset, we first extracted the genes involved in the iGenSig signature and then classified these genes into positive contributing genes and negative contributing genes. The positive contributing genes are defined as upregulated genes or genes with hotspot mutations. The negative contributing genes are defined as downregulated genes or mutated genes without mutation hotspots. The pathways enriched in the positive or negative contributing genes for Erlotinib resistance are analyzed by the Concept Signature Enrichment Analysis (CSEA) developed in our previous study^8^. The resulting top 30 pathways are disambiguated via correcting the crosstalk effects between pathways, to reveal independent pathway modules^28^. A p-value <0.01 is used as cutoff for disambiguation. The functional associations between the significant pathways are then assessed using our CSEA method as we previously described^8^, and the CSEA scores are then scaled between -1 and 1 and visualized using correlogram. The pathway network was visualized using the ‘igraph’ R package (ver. 1.2.4.2).

## Code availability

The R modules for iGenSig modeling are available through: https://github.com/wangxlab/iGenSig/.

## Acknowledgements

This study was supported by CCSG Dev Funds (P30 CA047904-31, X-S. W.), NIH grant 1R01CA181368 (X-S.W.), 1R01CA183976 (X-S.W.), 1R21CA237964 (X-S.W.), Commonwealth of PA Tobacco Phase 15 Formula Fund (X-S. W.), the Shear Family Foundation, and the Hillman Foundation. The results published here are in part based upon data generated by The Cancer Genome Atlas project established by the NCI and NHGRI (dbGaP accession: phs000178.v6.p6). This research was supported in part by the University of Pittsburgh Center for Research Computing through the resources provided. We especially thank Fangping Mu for the kind assistance of the research computing. This work also used the Extreme Science and Engineering Discovery Environment (XSEDE), which is supported by National Science Foundation grant number OCI-1053575. Specifically, it used the Bridges system, which is supported by NSF award number ACI-1445606, at the Pittsburgh Supercomputing Center (PSC).

## References

1. Schwartzberg, L., Kim, E.S., Liu, D. & Schrag, D. Precision Oncology: Who, How, What, When, and When Not? Am Soc Clin Oncol Educ Book 37, 160–169 (2017).

2. Zachary D. Stephens, S.Y.L., Faraz Faghri, Roy H. Campbell, Chengxiang Zhai, Miles J. Efron, Ravishankar Iyer, Michael C. Schatz, Saurabh Sinha, Gene E. Robinson. Challenges and Cases of Genomic Data Integration Across Technologies and Biological Scales.. (2018).

3. Vasconcellos, V.F., Colli, L.M., Awada, A. & de Castro Junior, G. Precision oncology: as much expectations as limitations. Ecancermedicalscience 12, ed86 (2018).

4. Frohlich, H. et al. From hype to reality: data science enabling personalized medicine. BMC Med 16, 150 (2018).

5. Yu, L., Zhou, D., Gao, L. & Zha, Y. Prediction of drug response in multilayer networks based on fusion of multiomics data. Methods (2020).

6. Jang, I.S., Neto, E.C., Guinney, J., Friend, S.H. & Margolin, A.A. Systematic assessment of analytical methods for drug sensitivity prediction from cancer cell line data. Pac Symp Biocomput, 63–74 (2014).

7. Liberzon, A. A description of the Molecular Signatures Database (MSigDB) Web site. Methods Mol Biol 1150, 153–160 (2014).

8. Chi, X. et al. Universal concept signature analysis: genome-wide quantification of new biological and pathological functions of genes and pathways. Brief Bioinform (2019).

9. Subramanian, A. et al. Gene set enrichment analysis: a knowledge-based approach for interpreting genome-wide expression profiles. Proc Natl Acad Sci U S A 102, 15545–15550 (2005).

10. Barretina, J. et al. The Cancer Cell Line Encyclopedia enables predictive modelling of anticancer drug sensitivity. Nature 483, 603–607 (2012).

11. Safikhani, Z. et al. Revisiting inconsistency in large pharmacogenomic studies. F1000Res 5, 2333 (2016).

12. Ding, M.Q., Chen, L., Cooper, G.F., Young, J.D. & Lu, X. Precision Oncology beyond Targeted Therapy: Combining Omics Data with Machine Learning Matches the Majority of Cancer Cells to Effective Therapeutics. Mol Cancer Res 16, 269–278 (2018).

13. Sharifi-Noghabi, H., Zolotareva, O., Collins, C.C. & Ester, M. MOLI: multi-omics late integration with deep neural networks for drug response prediction. Bioinformatics 35, i501–i509 (2019).

14. Gao, H. et al. High-throughput screening using patient-derived tumor xenografts to predict clinical trial drug response. Nature medicine 21, 1318–1325 (2015).

15. Byers, L.A. et al. An epithelial-mesenchymal transition gene signature predicts resistance to EGFR and PI3K inhibitors and identifies Axl as a therapeutic target for overcoming EGFR inhibitor resistance. Clin Cancer Res 19, 279–290 (2013).

16. Karachaliou, N. et al. BIM and mTOR expression levels predict outcome to erlotinib in EGFR-mutant non-small-cell lung cancer. Sci Rep 5, 17499 (2015).

17. Yu, H.A. et al. Concurrent Alterations in EGFR-Mutant Lung Cancers Associated with Resistance to EGFR Kinase Inhibitors and Characterization of MTOR as a Mediator of Resistance. Clin Cancer Res 24, 3108–3118 (2018).

18. Taube, J.H. et al. Core epithelial-to-mesenchymal transition interactome gene-expression signature is associated with claudin-low and metaplastic breast cancer subtypes. Proc Natl Acad Sci U S A 107, 15449–15454 (2010).

19. Yoshida, T. et al. ZEB1 Mediates Acquired Resistance to the Epidermal Growth Factor Receptor-Tyrosine Kinase Inhibitors in Non-Small Cell Lung Cancer. PLoS One 11, e0147344 (2016).

20. Tulchinsky, E., Demidov, O., Kriajevska, M., Barlev, N.A. & Imyanitov, E. EMT: A mechanism for escape from EGFR-targeted therapy in lung cancer. Biochim Biophys Acta Rev Cancer 1871, 29–39 (2019).

21. Zhang, P., Sun, Y. & Ma, L. ZEB1: at the crossroads of epithelial-mesenchymal transition, metastasis and therapy resistance. Cell Cycle 14, 481–487 (2015).

22. Horwitz, E. et al. Human and mouse VEGFA-amplified hepatocellular carcinomas are highly sensitive to sorafenib treatment. Cancer Discov 4, 730–743 (2014).

23. Martin, D., Galisteo, R. & Gutkind, J.S. CXCL8/IL8 stimulates vascular endothelial growth factor (VEGF) expression and the autocrine activation of VEGFR2 in endothelial cells by activating NFkappaB through the CBM (Carma3/Bcl10/Malt1) complex. J Biol Chem 284, 6038–6042 (2009).

24. Irizarry, R.A. et al. Exploration, normalization, and summaries of high density oligonucleotide array probe level data. Biostatistics 4, 249–264 (2003).

25. Iorio, F. et al. A landscape of pharmacogenomic interactions in cancer. Cell 166, 740–754 (2016).

26. Gulli, A. & Pal, S. Deep learning with Keras. (Packt Publishing Ltd, 2017).

27. Friedman, J., Hastie, T. & Tibshirani, R. Regularization paths for generalized linear models via coordinate descent. Journal of statistical software 33, 1 (2010).

28. Donato, M. et al. Analysis and correction of crosstalk effects in pathway analysis. Genome Res 23, 1885–1893 (2013).

29. Hilsenbeck, S.G. & Clark, G.M. Practical p-value adjustment for optimally selected cutpoints. Statistics in medicine 15, 103–112 (1996).

