## Supplementary Figures and Tables for "An integral genomic signature approach for tailored cancer targeted therapy using genome-wide sequencing data"

**SUPPLEMENTARY MATERIALS**

**Supplementary Figures**

High dimensional genomic features

Low dimensional principal components

Pathway or network modules

Matrix factorization

Pathway or network analysis

Deep learning

(i.e. Autoencoder)

Low dimensional representations

Dimensionality reduction

Synthetic features

Integral Genomic Signature Modeling (iGenSig)

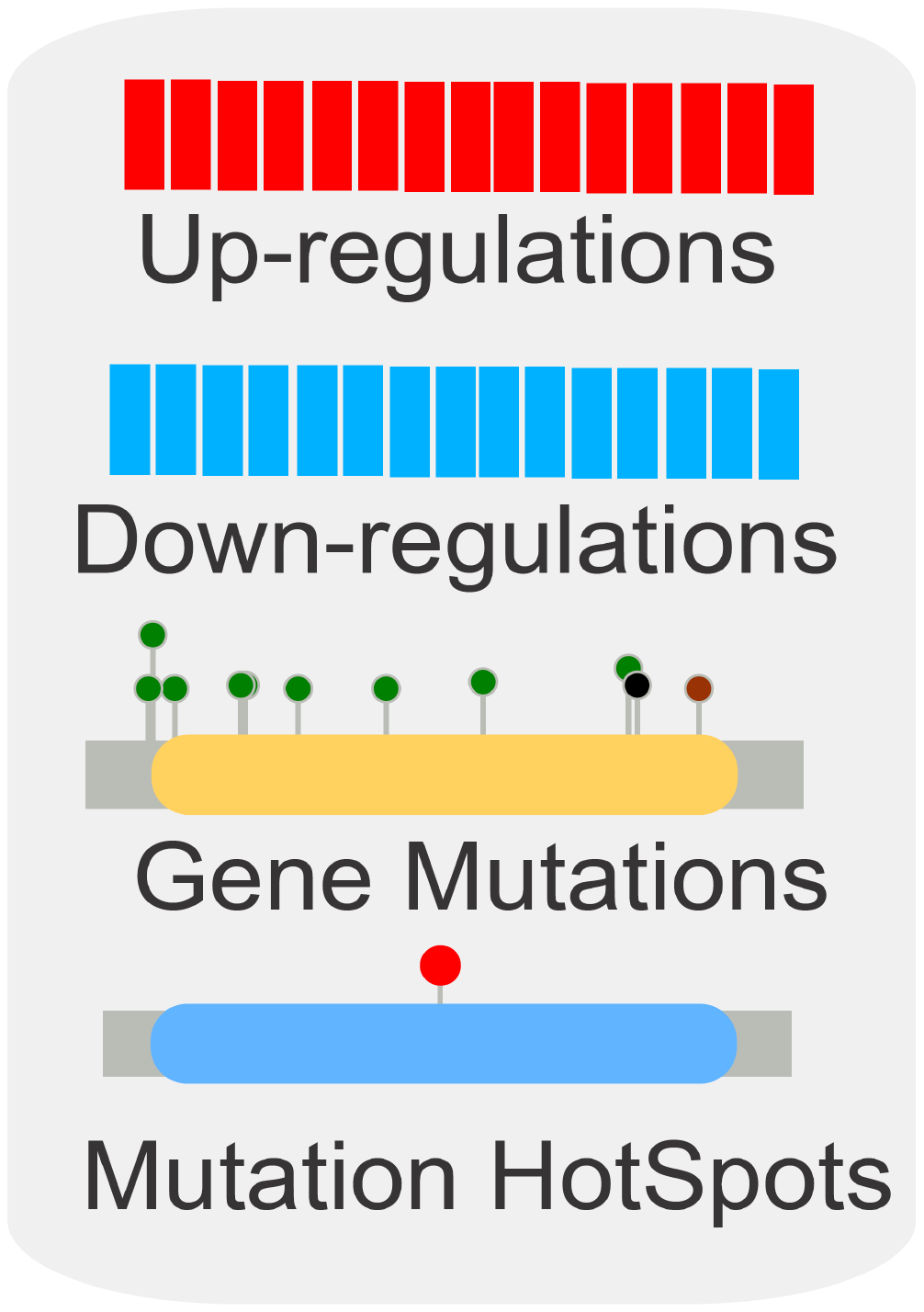

Feature removal

Gene signature panel

**Supplementary Figure 1. Schematic showing the key difference between integral genomic signature modeling and conventional gene signature or machine learning and deep learning methods in handling high-dimensional features.** An integral genomic signature is defined as the comprehensive set of high-dimensional genomic features predictive of a given clinical phenotype such as therapeutic response. iGenSig represents a new class of modeling methods that directly utilize high-dimensional genomic signature for predictive modeling based on multi-omics data.

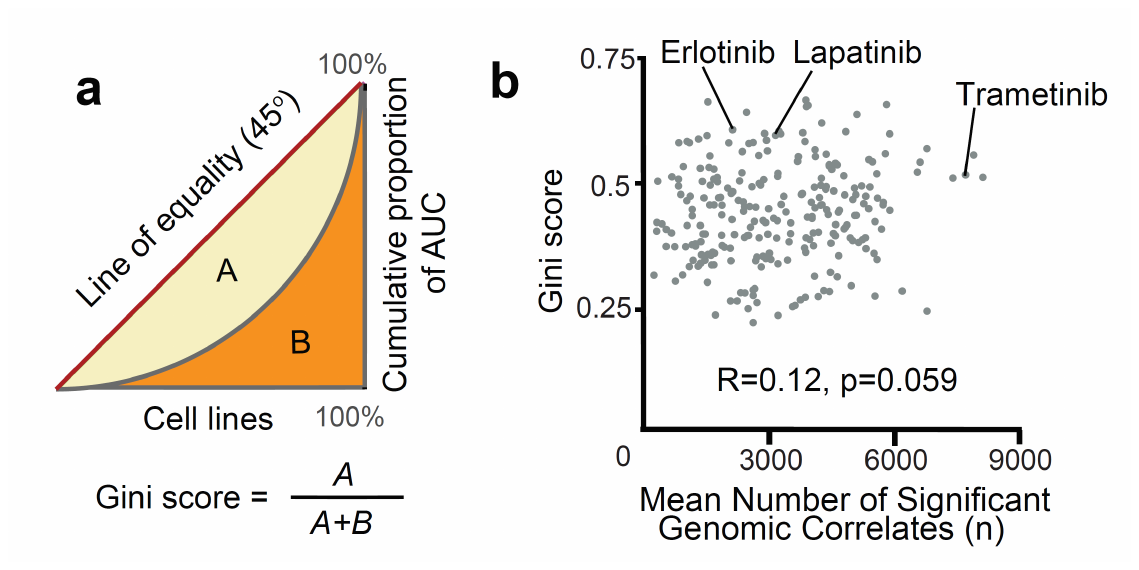

**Supplementary Figure 2. The GINI scores for the drugs profiled by GDSC do not correlate with their mean number of significant genomic correlates. a**) schematic showing the algorithm for calculating GINI scores for each drug based on unequal distribution of Act Area (1-AUC) drug sensitivity measurements among GDSC cell lines. **b**) Correlation between GINI scores and mean number of significant genomic correlates for the drugs profiled by GDSC.

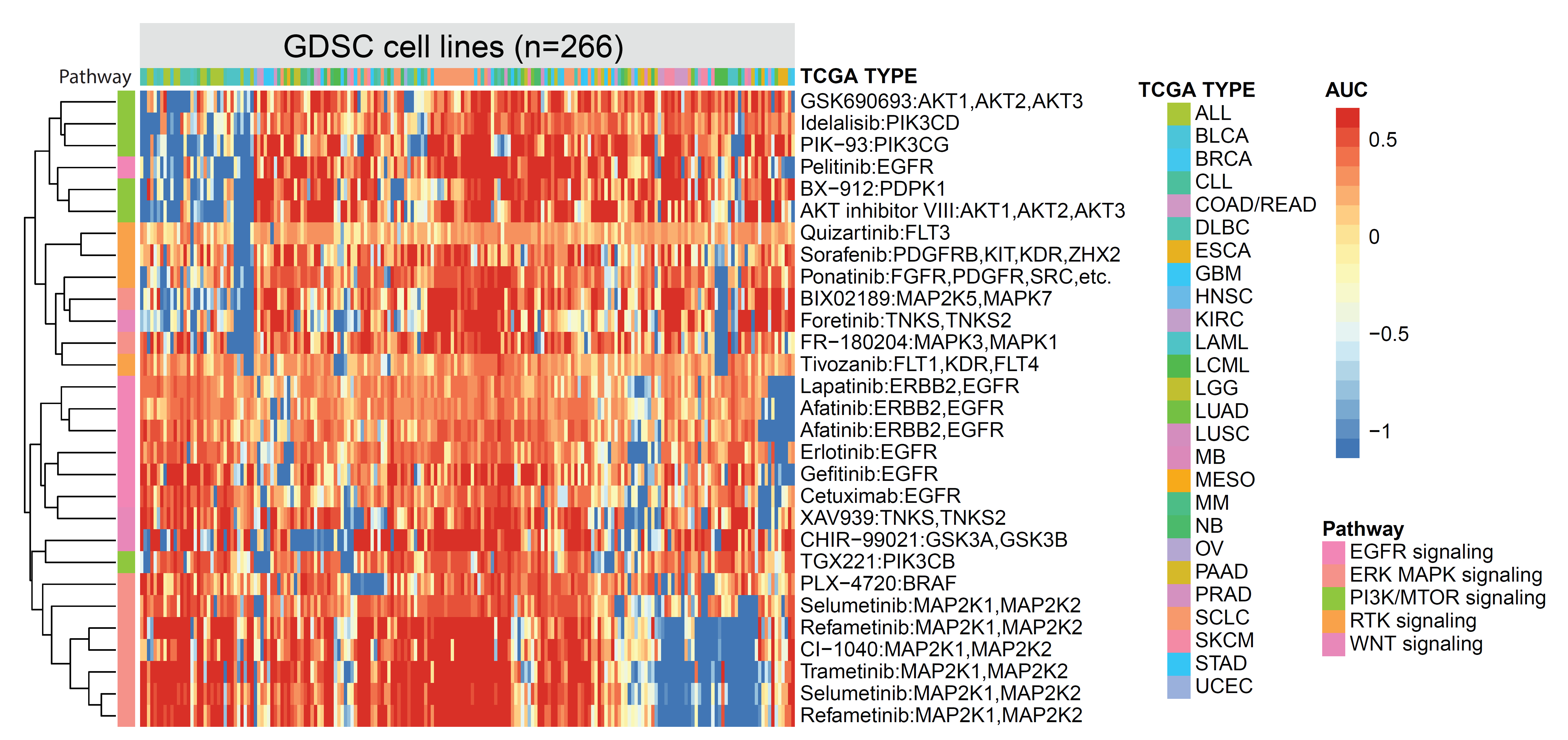

**Supplementary Figure 3. Clustering GDSC kinase-targeted drugs based on their AUC measurements.** The GDSC cell lines (n=266) with available AUC measurements for all the selected kinases are shown in the figure and are classified based on their TCGA cancer types. The pathways for the kinase targets are shown on the left bar.

Multi-omics features

Autoencoder (AE)

Elastic Net (EN)

Neural Net (NN)

Random Forest (RF)

Support Vector Machine (SVM)

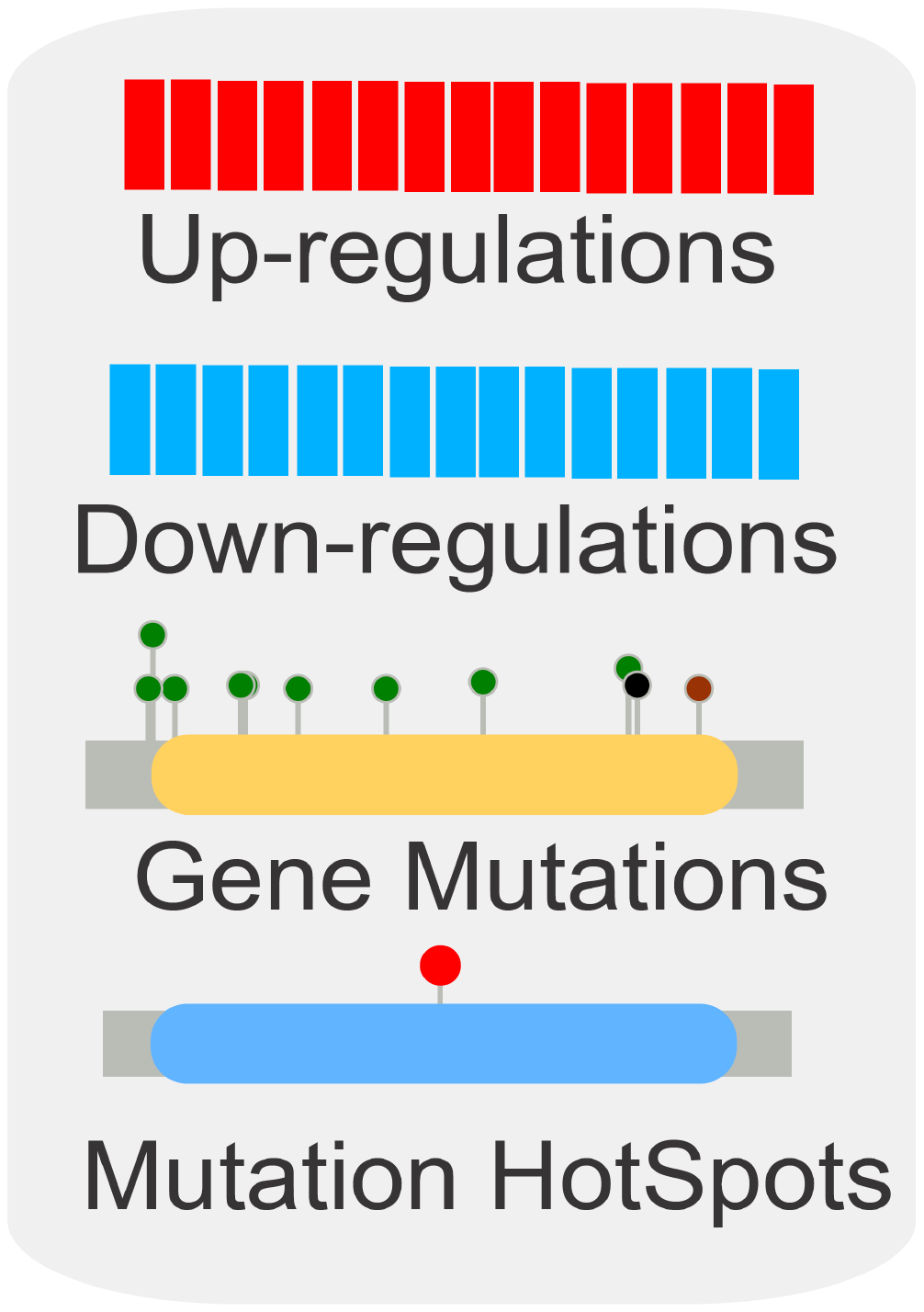

Unsupervised deep learning for dimension reduction

Supervised machine learning for sensitivity prediction

**Supplementary Figure 4. Schematic showing the workflow of deep learning and machine learning methods implemented in this study for drug sensitivity prediction.**

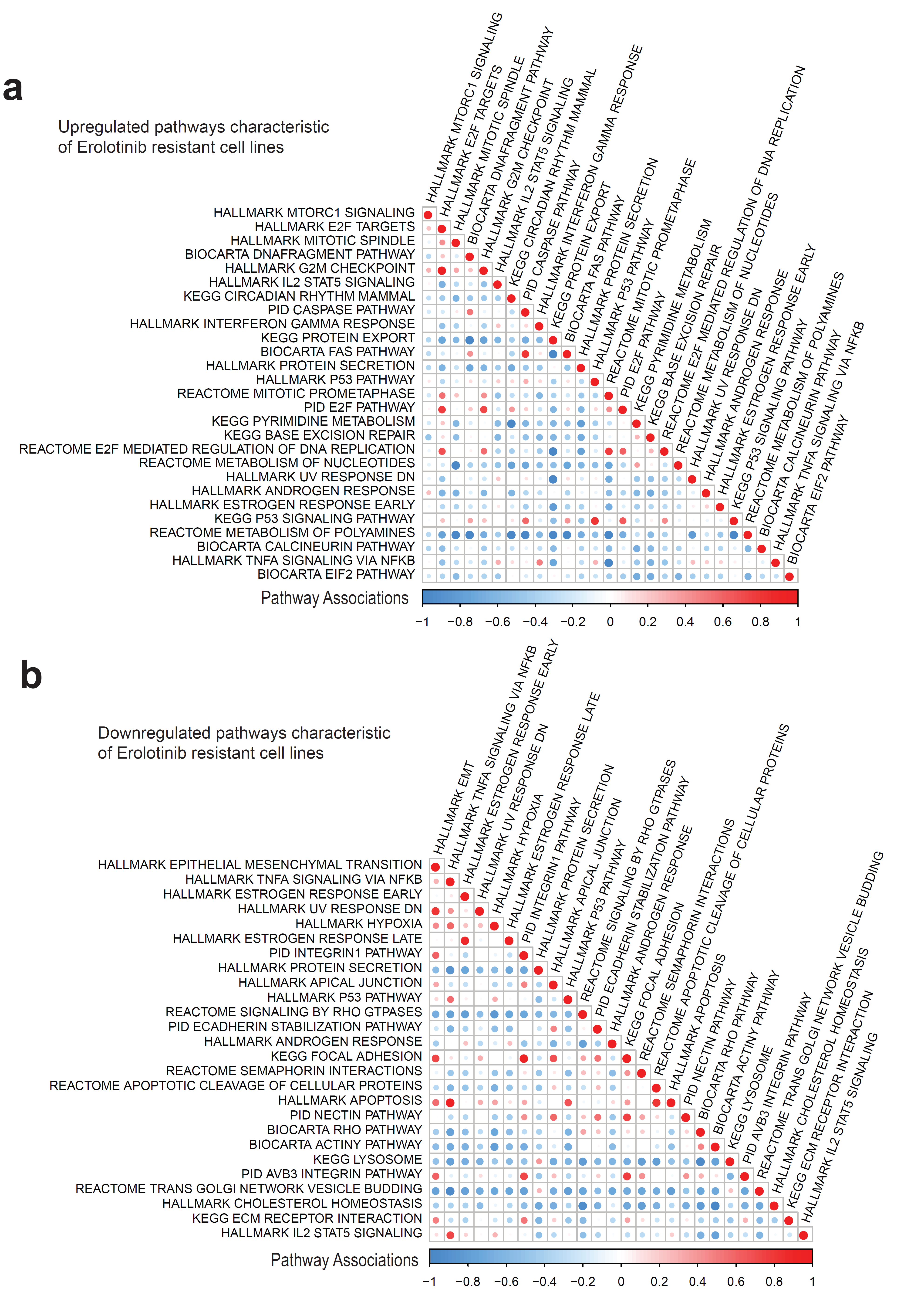

**Supplementary Figure 5. The upregulated or downregulated pathways characteristic of Erlotinib resistant iGenSig signature. a**) The upregulated pathways characteristic of Erlotinib resistant cell lines. **b**) the downregulated pathways characteristic of Erlotinib resistant cell lines. The function associations between the significant pathways assessed by concept signature enrichment analysis are shown in red to blue color scales. The pathways are sorted in descending order based their normalized enrichments scores with the most enriched pathways ranked on the top.

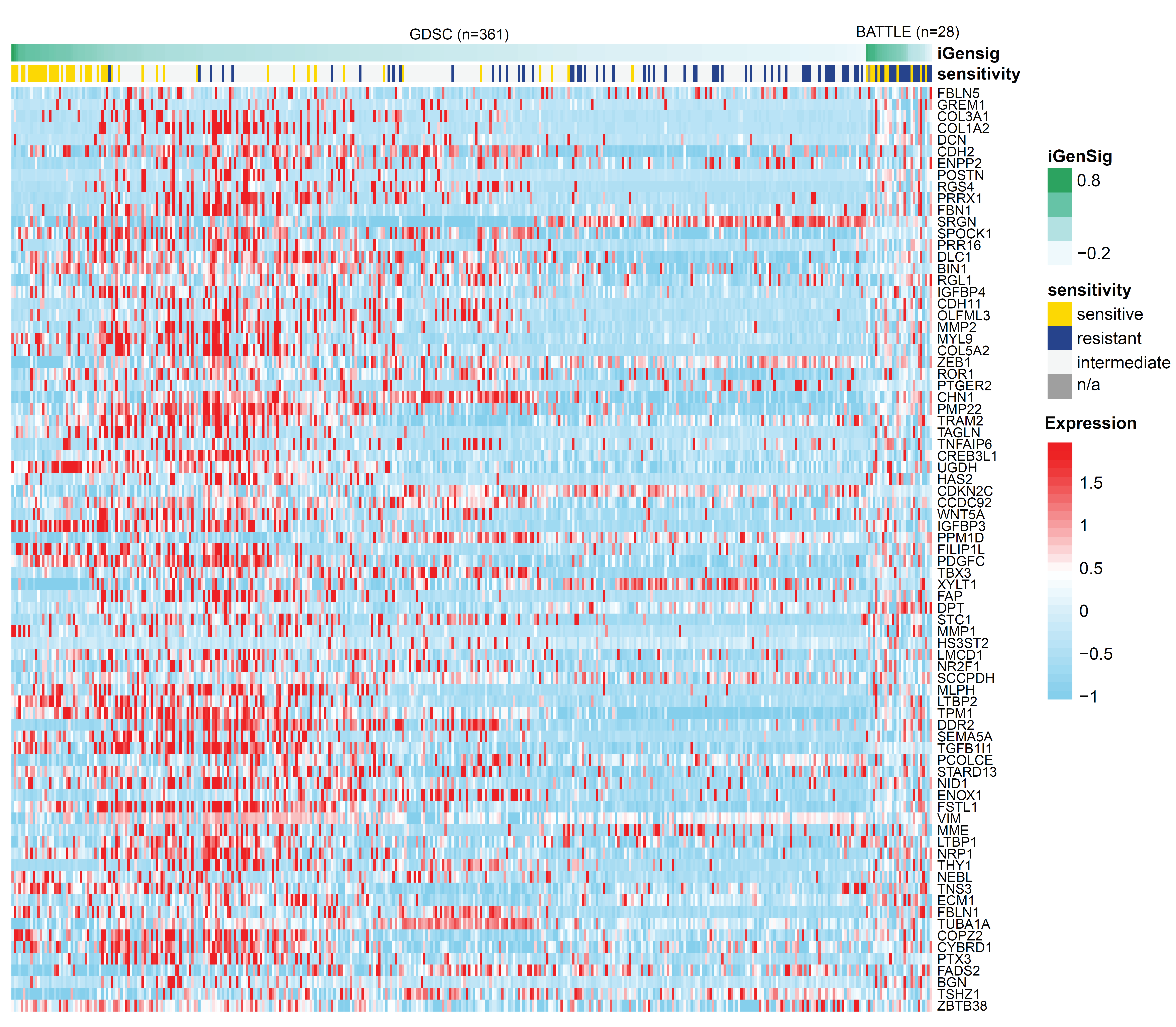

**Supplementary Figure 6. The expression of upregulated gene signature in EMT in association with the iGenSig scores for Erlotinib in GDSC cell lines and patient subjects from the BATTLE trial.** The cell line and patient subjects are sorted decreasingly by their iGenSig scores.

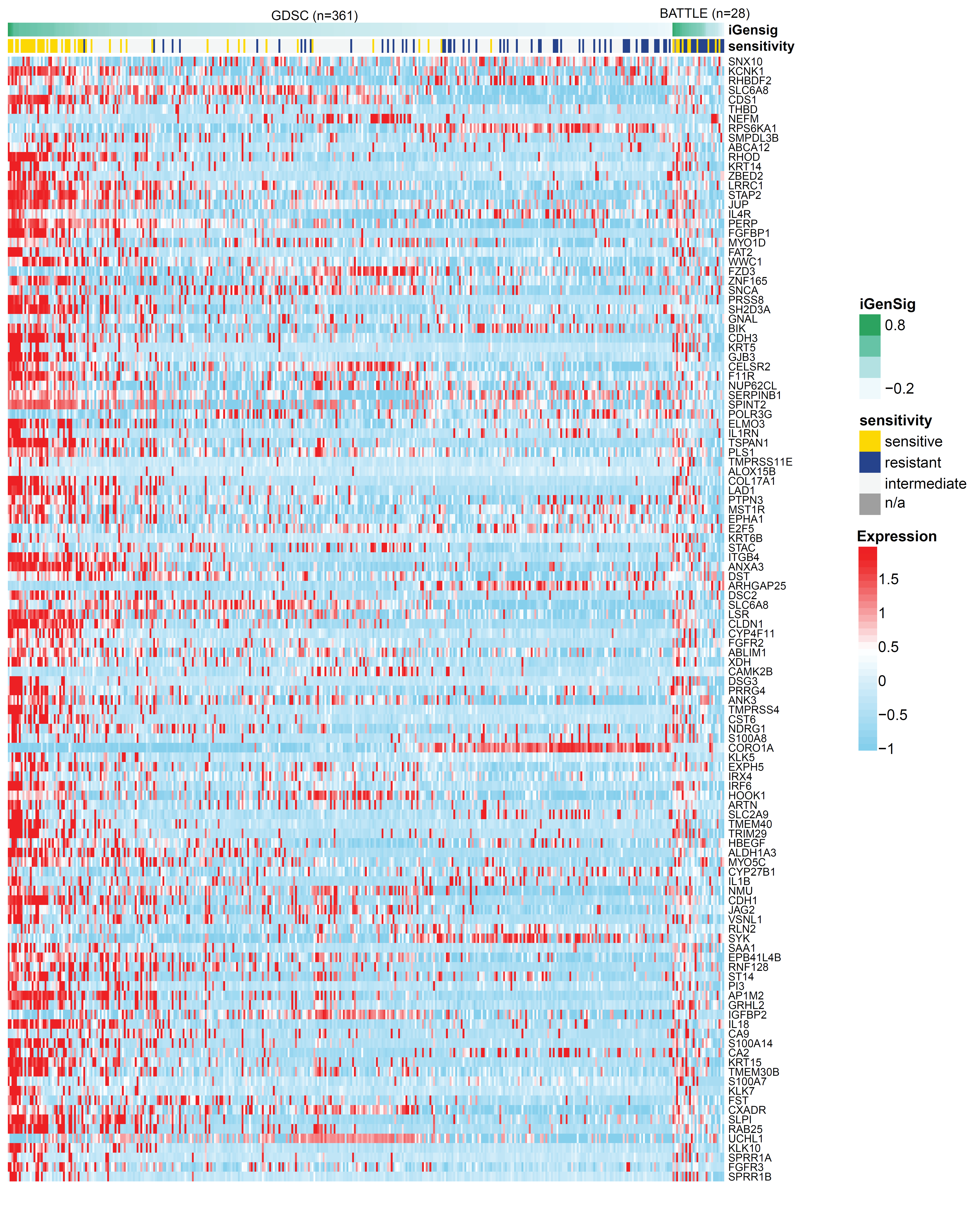

**Supplementary Figure 7. The expression of downregulated gene signature in EMT in association with the iGenSig scores for Erlotinib in GDSC cell lines and patient subjects from the BATTLE trial.** The cell line and patient subjects are sorted decreasingly by their iGenSig scores.

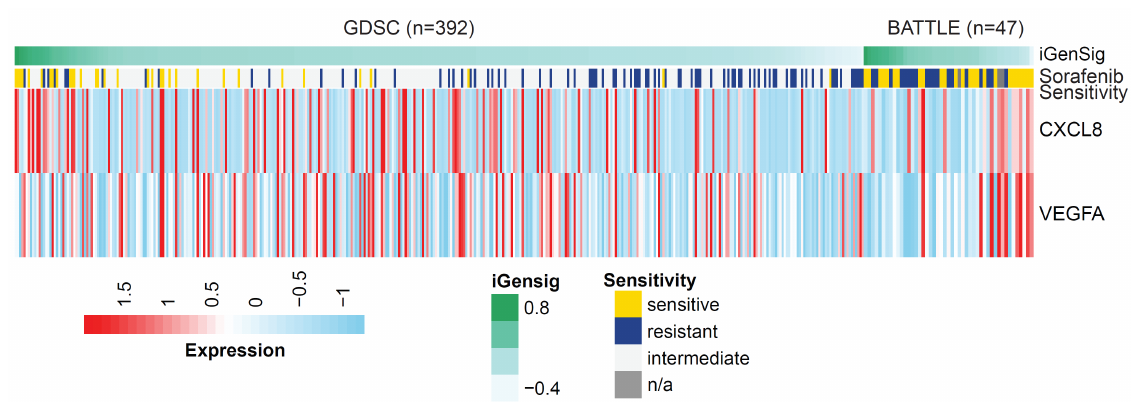

**Supplementary Figure 8. The expression of VEGFA and CXCL8 in correlation with the iGenSig scores for Erlotinib in GDSC cell lines and patient subjects from BATTLE trial.** The cell line and patient subjects are sorted decreasingly by their iGenSig scores.

**Supplementary Tables**

**Supplementary Table 1. A summary of the Pharmacogenomic and clinical trial datasets used in this study.**

| Genomic Dataset for Drug Sensitivity (GDSC) | Pharmacologic profiles for 267 anticancer drugs across 989 cell lines. |
| --- | --- |
|  | Affymetrix Human Genome U219 Array data |
|  | Whole exome sequencing data |
| Cancer Cell Line Encyclopedia (CCLE) | Pharmacologic profiles for 24 anticancer drugs across 504 cell lines. |
|  | Transcriptome sequencing data |
|  | Whole exome sequencing data |
| Pharmacogenomics Data for Patient Derived Xenografts | 1,000 patient-derived tumor xenograft models (PDXs) |
|  | 62 treatments across six indications |
|  | Transcriptome sequencing data |
|  | Whole exome sequencing data |
| Biomarker-integrated Approaches of Targeted Therapy for Lung Cancer Elimination (BATTLE). | 255 chemorefractory NSCLC patients were adaptively randomized to 4 arms: 1) erlotinib, 2) vandetanib, 3) erlotinib plus bexarotene, or 4) sorafenib. |
|  | Affymetrix Human Gene 1.0 ST Array data are available for 131 patients including patients treated with the following drugs profiled by GDSC: 1) Erolotinib (n=28), 2) Sorafanib (n=47). |

**Supplementary Table 2. Summary of iGenSig modeling results based on training and intenal testing sets using GDSC dataset.** The drugs with good performing iGenSig models are highlighted in grey (AUROC>0.75).

| drug ID | drugName | Target | Target Pathway | Sensitive Cell Line(n) | Resistant Cell Line(n) | Skewness | Kurtness | GINI Score | Cutoff Sensitive | Cutoff Resistant | Average AUROC | Stdv AUROC | Ave. num. of sig. features |
| --- | --- | --- | --- | --- | --- | --- | --- | --- | --- | --- | --- | --- | --- |
| 119 | Lapatinib | ERBB2, EGFR | EGFR signaling | 31 | 71 | -4.93 | 31.76 | 0.598 | 0.896 | 0.990 | **0.951** | 0.024 | 3372 |
| 276 | CAY10603 | HDAC1, HDAC6 | Chromatin histone acetylation | 129 | 226 | -0.01 | 2.48 | 0.285 | 0.335 | 0.727 | **0.911** | 0.032 | 5859 |
| 272 | AR-42 | HDAC1 | Chromatin histone acetylation | 171 | 228 | -0.02 | 2.34 | 0.295 | 0.395 | 0.756 | **0.902** | 0.029 | 6566 |
| 219 | AT-7519 | CDK1, CDK2, CDK4, CDK6, CDK9 | Cell cycle | 148 | 209 | -0.49 | 2.15 | 0.412 | 0.426 | 0.904 | **0.889** | 0.033 | 6030 |
| 260 | NG-25 | TAK1, MAP4K2 | Other, kinases | 108 | 249 | -2.84 | 12.93 | 0.485 | 0.738 | 0.954 | **0.877** | 0.027 | 4792 |
| 329 | QL-XI-92 | DDR1 | Other | 102 | 139 | -2.60 | 12.24 | 0.513 | 0.848 | 0.986 | **0.875** | 0.016 | 5051 |
| 333 | T0901317 | LXR, FXR | Other | 49 | 172 | -9.03 | 99.78 | 0.459 | 0.912 | 0.985 | **0.874** | 0.025 | 3426 |
| 152 | CP466722 | ATM | Genome integrity | 134 | 225 | -0.87 | 3.76 | 0.375 | 0.684 | 0.918 | **0.872** | 0.052 | 5628 |
| 257 | NPK76-II-72-1 | PLK3 | Cell cycle | 189 | 180 | -1.61 | 5.24 | 0.520 | 0.771 | 0.976 | **0.866** | 0.048 | 6959 |
| 1032 | Afatinib | ERBB2, EGFR | EGFR signaling | 146 | 114 | -3.18 | 14.40 | 0.649 | 0.893 | 0.989 | **0.861** | 0.037 | 6161 |
| 252 | WZ3105 | SRC, ROCK2, NTRK2, FLT3, IRAK1, others | Other | 174 | 214 | -1.22 | 4.76 | 0.463 | 0.659 | 0.951 | **0.854** | 0.037 | 6078 |
| 290 | KIN001-260 | IKKB | Other | 71 | 165 | -4.06 | 33.68 | 0.494 | 0.858 | 0.985 | **0.849** | 0.045 | 5519 |
| 295 | NVP-BHG712 | EPHB4 | Other | 163 | 194 | -2.90 | 13.32 | 0.534 | 0.812 | 0.974 | **0.846** | 0.018 | 4677 |
| 309 | Y-39983 | ROCK | Cytoskeleton | 104 | 141 | -3.31 | 17.44 | 0.526 | 0.843 | 0.983 | **0.841** | 0.022 | 4741 |
| 1377 | Afatinib | ERBB2, EGFR | EGFR signaling | 145 | 134 | -2.90 | 11.60 | 0.630 | 0.844 | 0.986 | **0.839** | 0.045 | 5395 |
| 273 | CUDC-101 | HDAC1-10, EGFR, ERBB2 | Other | 136 | 244 | -0.13 | 2.41 | 0.305 | 0.370 | 0.753 | **0.834** | 0.029 | 5247 |
| 261 | TL-1-85 | TAK | Other, kinases | 134 | 175 | -3.61 | 19.54 | 0.534 | 0.818 | 0.979 | **0.834** | 0.040 | 4882 |
| 275 | I-BET-762 | BRD2, BRD3, BRD4 | Chromatin other | 234 | 154 | -1.72 | 5.46 | 0.555 | 0.825 | 0.982 | **0.833** | 0.020 | 6130 |
| 94 | TGX221 | PI3Kbeta | PI3K/MTOR signaling | 50 | 50 | -4.01 | 25.48 | 0.576 | 0.886 | 0.990 | **0.829** | 0.044 | 2706 |
| 256 | JW-7-24-1 | LCK | Other, kinases | 243 | 200 | -0.85 | 3.17 | 0.366 | 0.669 | 0.884 | **0.827** | 0.027 | 5834 |
| 226 | GSK1070916 | AURKA, AURKC | Mitosis | 156 | 156 | -2.50 | 8.37 | 0.657 | 0.811 | 0.987 | **0.824** | 0.027 | 4068 |
| 1011 | Navitoclax | BCL2, BCL-XL, BCL-W | Apoptosis regulation | 250 | 132 | -1.86 | 5.99 | 0.593 | 0.855 | 0.981 | **0.820** | 0.024 | 6243 |
| 271 | VNLG/124 | HDAC,RAR | Chromatin histone acetylation | 67 | 171 | -9.47 | 131.93 | 0.459 | 0.936 | 0.987 | **0.818** | 0.053 | 3435 |
| 155 | Ponatinib | ABL, PDGFRA, VEGFR2, FGFR1, SRC, TIE2, FLT3 | RTK signaling | 88 | 187 | -3.91 | 20.79 | 0.578 | 0.804 | 0.983 | **0.817** | 0.062 | 4044 |
| 1010 | Gefitinib | EGFR | EGFR signaling | 84 | 132 | -3.97 | 23.89 | 0.537 | 0.888 | 0.987 | **0.816** | 0.057 | 4834 |
| 1114 | Cetuximab | EGFR | EGFR signaling | 86 | 158 | -5.45 | 43.97 | 0.539 | 0.906 | 0.987 | **0.816** | 0.033 | 7031 |
| 228 | AKT inhibitor VIII | AKT1, AKT2, AKT3 | PI3K/MTOR signaling | 184 | 190 | -1.61 | 5.54 | 0.468 | 0.759 | 0.961 | **0.815** | 0.040 | 5287 |
| 345 | KIN001-270 | CDK9 | Cell cycle | 63 | 157 | -3.55 | 28.07 | 0.376 | 0.915 | 0.984 | **0.815** | 0.052 | 4034 |
| 1037 | BX795 | TBK1, PDK1 (PDPK1), IKK, AURKB, AURKC | Other | 66 | 204 | -1.79 | 10.31 | 0.368 | 0.731 | 0.935 | **0.813** | 0.031 | 4064 |
| 265 | Tubastatin A | HDAC1, HDAC6, HDAC8 | Chromatin histone acetylation | 189 | 131 | -2.06 | 7.98 | 0.517 | 0.885 | 0.985 | **0.811** | 0.036 | 5906 |
| 306 | Fedratinib | JAK2 | Other, kinases | 178 | 209 | -1.40 | 5.21 | 0.438 | 0.713 | 0.943 | **0.810** | 0.040 | 5337 |
| 279 | BIX02189 | MEK5, ERK5 | ERK MAPK signaling | 94 | 182 | -3.31 | 20.51 | 0.477 | 0.847 | 0.980 | **0.810** | 0.072 | 5075 |
| 1014 | Refametinib | MEK1, MEK2 | ERK MAPK signaling | 186 | 171 | -1.35 | 4.52 | 0.509 | 0.694 | 0.971 | **0.809** | 0.033 | 7880 |
| 238 | Idelalisib | PI3Kdelta | PI3K/MTOR signaling | 93 | 165 | -4.15 | 27.38 | 0.541 | 0.848 | 0.984 | **0.808** | 0.061 | 3857 |
| 331 | QL-X-138 | BTK | Other, kinases | 228 | 195 | -0.71 | 2.78 | 0.400 | 0.612 | 0.907 | **0.803** | 0.025 | 4834 |
| 286 | KIN001-236 | Angiopoietin-1 receptor | Other | 88 | 169 | -2.78 | 14.86 | 0.416 | 0.919 | 0.985 | **0.803** | 0.033 | 5094 |
| 303 | PIK-93 | PI3Kgamma | PI3K/MTOR signaling | 188 | 201 | -1.50 | 4.98 | 0.451 | 0.678 | 0.936 | **0.800** | 0.031 | 5699 |
| 310 | YM201636 | PYKFYVE | Other | 137 | 206 | -1.52 | 6.20 | 0.420 | 0.804 | 0.961 | **0.800** | 0.059 | 5133 |
| 1008 | Methotrexate | Antimetabolite | DNA replication | 180 | 91 | -2.00 | 6.62 | 0.597 | 0.835 | 0.986 | **0.798** | 0.057 | 5103 |
| 287 | KIN001-244 | PDK1 (PDPK1) | Other, kinases | 125 | 219 | -2.40 | 11.74 | 0.393 | 0.774 | 0.937 | **0.797** | 0.031 | 4169 |
| 245 | UNC0638 | G9a and GLP methyltransferases | Chromatin histone methylation | 137 | 236 | -0.31 | 2.35 | 0.355 | 0.637 | 0.893 | **0.797** | 0.048 | 5913 |
| 1001 | AICA Ribonucleotide | AMPK agonist | Metabolism | 100 | 187 | -1.33 | 5.29 | 0.396 | 0.726 | 0.945 | **0.796** | 0.048 | 5219 |
| 1372 | Trametinib | MEK1, MEK2 | ERK MAPK signaling | 382 | 159 | -0.79 | 2.33 | 0.515 | 0.722 | 0.973 | **0.795** | 0.024 | 8208 |
| 305 | TPCA-1 | IKK2 | Other, kinases | 203 | 203 | -1.50 | 5.14 | 0.471 | 0.730 | 0.951 | **0.795** | 0.038 | 5925 |
| 326 | GSK690693 | AKT1, AKT2, AKT3 | PI3K/MTOR signaling | 163 | 135 | -3.15 | 13.65 | 0.645 | 0.853 | 0.987 | **0.794** | 0.031 | 4082 |
| 1022 | AZD7762 | CHEK1, CHEK2 | Cell cycle | 63 | 218 | -0.72 | 3.91 | 0.322 | 0.467 | 0.837 | **0.793** | 0.046 | 4882 |
| 222 | BX-912 | PDK1 (PDPK1) | PI3K/MTOR signaling | 178 | 172 | -2.05 | 7.31 | 0.544 | 0.757 | 0.974 | **0.793** | 0.052 | 5704 |
| 301 | PHA-793887 | CDK2, CDK7, CDK5 | Cell cycle | 262 | 166 | -1.46 | 4.59 | 0.540 | 0.744 | 0.979 | **0.793** | 0.042 | 5793 |
| 1 | Erlotinib | EGFR | EGFR signaling | 49 | 61 | -4.25 | 24.56 | 0.601 | 0.922 | 0.989 | **0.792** | 0.078 | 2172 |
| 253 | XMD14-99 | ALK, CDK7, LTK, others | Other | 111 | 156 | -4.66 | 41.26 | 0.448 | 0.929 | 0.987 | **0.790** | 0.050 | 4937 |
| 56 | WH-4-023 | SRC, LCK | Other, kinases | 102 | 50 | -2.03 | 6.50 | 0.653 | 0.829 | 0.987 | **0.788** | 0.025 | 1529 |
| 1526 | Refametinib | MEK1, MEK2 | ERK MAPK signaling | 159 | 190 | -1.57 | 5.43 | 0.510 | 0.707 | 0.971 | **0.785** | 0.015 | 8656 |
| 1498 | Selumetinib | MEK1, MEK2 | ERK MAPK signaling | 211 | 163 | -1.75 | 5.71 | 0.552 | 0.765 | 0.979 | **0.783** | 0.015 | 8414 |
| 179 | 5-Fluorouracil | Antimetabolite (DNA & RNA) | Other | 234 | 183 | -1.36 | 4.55 | 0.495 | 0.742 | 0.966 | **0.781** | 0.040 | 4459 |
| 1062 | Selumetinib | MEK1, MEK2 | ERK MAPK signaling | 148 | 109 | -2.62 | 10.33 | 0.613 | 0.864 | 0.988 | **0.781** | 0.033 | 4482 |
| 1241 | CHIR-99021 | GSK3A, GSK3B | WNT signaling | 69 | 154 | -2.05 | 9.80 | 0.463 | 0.849 | 0.984 | **0.780** | 0.035 | 4210 |
| 59 | WZ-1-84 | BMX | Other, kinases | 59 | 66 | -2.81 | 13.02 | 0.507 | 0.903 | 0.986 | **0.778** | 0.051 | 1684 |
| 1371 | PLX-4720 | BRAF | ERK MAPK signaling | 104 | 139 | -3.93 | 21.52 | 0.549 | 0.897 | 0.986 | **0.778** | 0.075 | 3876 |
| 1242 | (5Z)-7-Oxozeaenol | TAK1 | Other, kinases | 159 | 203 | -1.07 | 4.43 | 0.296 | 0.530 | 0.806 | **0.777** | 0.055 | 4537 |
| 1007 | Docetaxel | Microtubule stabiliser | Mitosis | 179 | 190 | -0.64 | 2.78 | 0.390 | 0.652 | 0.919 | **0.777** | 0.041 | 5283 |
| 211 | TL-2-105 | not defined | Unclassified | 101 | 151 | -4.22 | 27.29 | 0.550 | 0.869 | 0.985 | **0.775** | 0.049 | 4317 |
| 312 | Tivozanib | VEGFR1, VEGFR2, VEGFR3 | RTK signaling | 62 | 203 | -9.01 | 110.72 | 0.400 | 0.960 | 0.988 | **0.775** | 0.044 | 2588 |
| 88 | Entinostat | HDAC1, HDAC3 | Chromatin histone acetylation | 49 | 106 | -0.45 | 2.76 | 0.348 | 0.538 | 0.867 | **0.774** | 0.081 | 1345 |
| 1012 | Vorinostat | HDAC inhibitor Class I, IIa, IIb, IV | Chromatin histone acetylation | 99 | 197 | -0.38 | 2.65 | 0.257 | 0.519 | 0.810 | **0.774** | 0.041 | 7212 |
| 292 | Masitinib | KIT, PDGFRA, PDGFRB | Other, kinases | 75 | 293 | -3.81 | 24.25 | 0.406 | 0.789 | 0.943 | **0.773** | 0.046 | 4397 |
| 308 | Foretinib | MET, KDR, TIE2, VEGFR3/FLT4, RON, PDGFR, FGFR1, EGFR | RTK signaling | 88 | 250 | -0.53 | 3.96 | 0.278 | 0.488 | 0.788 | **0.773** | 0.027 | 3931 |
| 1268 | XAV939 | TNKS1, TNKS2 | WNT signaling | 88 | 132 | -2.94 | 15.97 | 0.461 | 0.901 | 0.986 | **0.768** | 0.063 | 5570 |
| 30 | Sorafenib | PDGFR, KIT, VEGFR, RAF | RTK signaling | 33 | 96 | -4.53 | 30.07 | 0.450 | 0.804 | 0.964 | **0.768** | 0.082 | 1129 |
| 1024 | Lestaurtinib | FLT3, JAK2, NTRK1, NTRK2, NTRK3 | Other, kinases | 121 | 212 | -0.62 | 3.01 | 0.323 | 0.485 | 0.830 | **0.766** | 0.077 | 4294 |
| 294 | MPS-1-IN-1 | MPS1 | Mitosis | 86 | 196 | -2.06 | 10.81 | 0.449 | 0.775 | 0.974 | **0.766** | 0.089 | 4612 |
| 282 | Pelitinib | EGFR | EGFR signaling | 158 | 150 | -2.14 | 7.94 | 0.590 | 0.758 | 0.987 | **0.764** | 0.035 | 3283 |
| 203 | BMS-345541 | IKK1, IKK2 | Other, kinases | 135 | 231 | -1.72 | 7.37 | 0.362 | 0.793 | 0.929 | **0.762** | 0.052 | 4917 |
| 254 | Quizartinib | FLT3 | RTK signaling | 51 | 162 | -9.51 | 109.24 | 0.504 | 0.915 | 0.988 | **0.761** | 0.081 | 2350 |
| 263 | FR-180204 | ERK1, ERK2 | ERK MAPK signaling | 46 | 160 | -4.72 | 49.06 | 0.416 | 0.901 | 0.986 | **0.761** | 0.072 | 2635 |
| 1015 | CI-1040 | MEK1, MEK2 | ERK MAPK signaling | 165 | 169 | -1.67 | 5.98 | 0.448 | 0.732 | 0.945 | **0.759** | 0.046 | 6248 |
| 51 | Dasatinib | ABL, SRC, Ephrins, PDGFR, KIT | Other | 160 | 42 | -1.42 | 3.95 | 0.634 | 0.870 | 0.988 | **0.758** | 0.041 | 2533 |
| 330 | XMD13-2 | RIPK1 | Apoptosis regulation | 159 | 177 | -2.04 | 7.96 | 0.475 | 0.853 | 0.977 | **0.757** | 0.052 | 5622 |
| 55 | A-770041 | LCK, FYN | Other, kinases | 83 | 47 | -2.06 | 7.67 | 0.583 | 0.772 | 0.986 | **0.756** | 0.069 | 1292 |
| 300 | CX-5461 | RNA Polymerase 1 | Other | 212 | 133 | -1.84 | 6.06 | 0.595 | 0.775 | 0.986 | **0.755** | 0.055 | 3990 |
| 230 | GSK429286A | ROCK1, ROCK2 | Cytoskeleton | 92 | 140 | -4.62 | 32.43 | 0.518 | 0.892 | 0.986 | **0.753** | 0.057 | 3346 |
| 1026 | Tanespimycin | HSP90 | Protein stability and degradation | 166 | 170 | -0.72 | 2.59 | 0.458 | 0.577 | 0.951 | **0.751** | 0.014 | 6071 |
| 1378 | Bleomycin (50 uM) | dsDNA break induction | DNA replication | 314 | 213 | -0.56 | 2.25 | 0.404 | 0.546 | 0.877 | **0.750** | 0.031 | 4785 |
| 1061 | SB590885 | BRAF | ERK MAPK signaling | 75 | 132 | -5.58 | 41.40 | 0.563 | 0.915 | 0.988 | **0.747** | 0.101 | 4308 |
| 1047 | Nutlin-3a (-) | MDM2 | p53 pathway | 214 | 117 | -1.84 | 6.19 | 0.560 | 0.912 | 0.987 | **0.743** | 0.033 | 3075 |
| 332 | XMD15-27 | CAMK2 | Other, kinases | 81 | 170 | -4.25 | 29.34 | 0.403 | 0.944 | 0.988 | **0.741** | 0.103 | 3085 |
| 1373 | Dabrafenib | BRAF | ERK MAPK signaling | 129 | 136 | -2.99 | 11.75 | 0.647 | 0.809 | 0.984 | **0.741** | 0.061 | 4126 |
| 1495 | Olaparib | PARP1, PARP2 | Genome integrity | 85 | 174 | -3.05 | 19.81 | 0.433 | 0.866 | 0.979 | **0.741** | 0.043 | 3234 |
| 159 | HG6-64-1 | BRAF | ERK MAPK signaling | 100 | 212 | -1.04 | 4.38 | 0.404 | 0.558 | 0.912 | **0.740** | 0.068 | 3676 |
| 1494 | SN-38 | TOP1 | DNA replication | 137 | 223 | -0.37 | 2.24 | 0.330 | 0.423 | 0.829 | **0.738** | 0.034 | 4708 |
| 1060 | PD0325901 | MEK1, MEK2 | ERK MAPK signaling | 189 | 155 | -1.59 | 5.32 | 0.565 | 0.752 | 0.988 | **0.737** | 0.073 | 7209 |
| 344 | THZ-2-49 | CDK9 | Cell cycle | 381 | 144 | -0.73 | 2.44 | 0.509 | 0.740 | 0.984 | **0.737** | 0.026 | 4473 |
| 53 | CGP-60474 | CDK1,CDK2,CDK5,CDK7,CDK9, PKC | Cell cycle | 58 | 109 | -0.14 | 2.45 | 0.299 | 0.632 | 0.857 | **0.736** | 0.028 | 2746 |
| 106 | XMD8-85 | ERK5, BET | Other | 34 | 93 | -1.28 | 5.73 | 0.403 | 0.789 | 0.961 | **0.735** | 0.050 | 1462 |
| 1059 | AZD8055 | MTORC1, MTORC2 | PI3K/MTOR signaling | 143 | 208 | -0.93 | 3.99 | 0.294 | 0.666 | 0.862 | **0.734** | 0.019 | 4685 |
| 154 | CHIR-99021 | GSK3A, GSK3B | WNT signaling | 125 | 135 | -3.98 | 24.24 | 0.492 | 0.938 | 0.989 | **0.734** | 0.045 | 2674 |
| 1266 | ICL1100013 | N-myristoyltransferase 1/2 | Other | 119 | 181 | -0.65 | 2.56 | 0.385 | 0.429 | 0.893 | **0.733** | 0.037 | 4721 |
| 283 | Omipalisib | PI3K (class 1), MTORC1, MTORC2 | PI3K/MTOR signaling | 256 | 191 | -0.82 | 2.93 | 0.424 | 0.611 | 0.913 | **0.733** | 0.086 | 3680 |
| 52 | GNF-2 | BCR-ABL | ABL signaling | 29 | 78 | -5.70 | 40.10 | 0.506 | 0.942 | 0.989 | **0.732** | 0.128 | 2982 |
| 1013 | Nilotinib | ABL | ABL signaling | 45 | 128 | -6.77 | 50.92 | 0.593 | 0.902 | 0.988 | **0.732** | 0.072 | 3001 |
| 262 | VX-11e | ERK2 | ERK MAPK signaling | 155 | 133 | -2.55 | 11.39 | 0.559 | 0.844 | 0.985 | **0.731** | 0.036 | 2420 |
| 1259 | Talazoparib | PARP1, PARP2 | Genome integrity | 174 | 177 | -1.62 | 5.49 | 0.492 | 0.661 | 0.953 | **0.729** | 0.043 | 5728 |
| 288 | WHI-P97 | JAK3 | Other, kinases | 94 | 172 | -3.42 | 19.03 | 0.473 | 0.899 | 0.984 | **0.729** | 0.029 | 1861 |
| 196 | Phenformin | Biguanide agent | Other | 271 | 201 | -1.01 | 3.53 | 0.440 | 0.665 | 0.935 | **0.728** | 0.047 | 4653 |
| 206 | Ruxolitinib | JAK1, JAK2 | Other, kinases | 58 | 189 | -10.31 | 125.76 | 0.467 | 0.949 | 0.988 | **0.728** | 0.046 | 2320 |
| 1050 | ZM447439 | AURKA, AURKB | Mitosis | 146 | 165 | -1.35 | 4.79 | 0.432 | 0.843 | 0.971 | **0.728** | 0.039 | 4231 |
| 29 | AZ628 | BRAF | ERK MAPK signaling | 99 | 73 | -2.04 | 6.80 | 0.575 | 0.809 | 0.976 | **0.725** | 0.091 | 2594 |
| 258 | STF-62247 | Autophagy inducer | Other | 132 | 150 | -3.54 | 19.32 | 0.445 | 0.941 | 0.987 | **0.724** | 0.048 | 3556 |
| 304 | SB52334 | ALK5 | RTK signaling | 172 | 161 | -2.51 | 10.38 | 0.519 | 0.876 | 0.982 | **0.722** | 0.031 | 2413 |
| 225 | Genentech Cpd 10 | AURKA, AURKB | Mitosis | 252 | 226 | -1.40 | 4.64 | 0.488 | 0.776 | 0.962 | **0.722** | 0.030 | 4587 |
| 1004 | Vinblastine | Microtubule destabiliser | Mitosis | 100 | 223 | -0.05 | 2.38 | 0.285 | 0.414 | 0.765 | **0.721** | 0.049 | 4391 |
| 171 | AKT inhibitor VIII | AKT1, AKT2, AKT3 | PI3K/MTOR signaling | 78 | 176 | -2.23 | 10.42 | 0.451 | 0.864 | 0.982 | **0.720** | 0.043 | 3107 |
| 1203 | QL-XII-61 | BMX, BTK | Other, kinases | 77 | 76 | -2.47 | 10.73 | 0.510 | 0.881 | 0.984 | **0.720** | 0.077 | 1775 |
| 134 | Etoposide | TOP2 | DNA replication | 136 | 199 | -0.52 | 2.33 | 0.401 | 0.419 | 0.895 | **0.718** | 0.065 | 1890 |
| 224 | AS605240 | PI3Kgamma | PI3K/MTOR signaling | 171 | 201 | -1.85 | 6.86 | 0.499 | 0.711 | 0.960 | **0.717** | 0.056 | 2863 |
| 221 | TAK-715 | p38alpha, p38beta | JNK and p38 signaling | 122 | 202 | -2.40 | 11.48 | 0.427 | 0.869 | 0.976 | **0.717** | 0.040 | 5859 |
| 1006 | Cytarabine | Antimetabolite | DNA replication | 176 | 188 | -0.97 | 3.54 | 0.421 | 0.638 | 0.924 | **0.715** | 0.026 | 4842 |
| 235 | QL-XII-47 | BTK, BMX | Other, kinases | 141 | 211 | -0.42 | 2.23 | 0.396 | 0.501 | 0.905 | **0.715** | 0.028 | 2862 |
| 1161 | ZG-10 | JNK1 | JNK and p38 signaling | 48 | 129 | -0.24 | 3.14 | 0.273 | 0.603 | 0.836 | **0.714** | 0.046 | 2806 |
| 176 | IPA-3 | PAK1 | Cytoskeleton | 252 | 214 | -0.95 | 3.28 | 0.478 | 0.814 | 0.976 | **0.712** | 0.015 | 3958 |
| 1175 | Rucaparib | PARP1, PARP2 | Genome integrity | 96 | 155 | -3.22 | 19.63 | 0.453 | 0.914 | 0.987 | **0.712** | 0.072 | 4306 |
| 38 | Saracatinib | ABL, SRC | RTK signaling | 63 | 55 | -2.94 | 14.17 | 0.576 | 0.883 | 0.989 | **0.711** | 0.097 | 2077 |
| 1009 | Tretinoin | Retinoic acid | Other | 75 | 134 | -5.37 | 39.25 | 0.559 | 0.871 | 0.986 | **0.710** | 0.048 | 2425 |
| 208 | Ispinesib Mesylate | KSP | Mitosis | 366 | 146 | -0.91 | 2.84 | 0.493 | 0.825 | 0.977 | **0.710** | 0.046 | 4601 |
| 165 | DMOG | HIF-PH | Metabolism | 113 | 214 | -0.30 | 2.43 | 0.283 | 0.376 | 0.776 | **0.709** | 0.101 | 4054 |
| 1030 | KU-55933 | ATM | Genome integrity | 44 | 169 | -3.03 | 21.39 | 0.400 | 0.880 | 0.981 | **0.709** | 0.085 | 2550 |
| 223 | ZSTK474 | PI3K (class 1) | PI3K/MTOR signaling | 225 | 206 | -1.17 | 4.02 | 0.435 | 0.653 | 0.924 | **0.708** | 0.034 | 4108 |
| 1019 | Bosutinib | SRC, ABL, TEC | Other, kinases | 158 | 146 | -1.78 | 6.47 | 0.512 | 0.843 | 0.983 | **0.708** | 0.067 | 3338 |
| 281 | Alectinib | ALK | RTK signaling | 58 | 172 | -7.72 | 74.63 | 0.551 | 0.890 | 0.987 | **0.705** | 0.071 | 1583 |
| 231 | FMK | RSK | Other, kinases | 57 | 136 | -4.57 | 37.01 | 0.402 | 0.931 | 0.987 | **0.704** | 0.099 | 2221 |
| 64 | CMK | RSK2 | ERK MAPK signaling | 32 | 85 | -0.98 | 3.81 | 0.425 | 0.781 | 0.973 | **0.703** | 0.067 | 213 |
| 277 | Linifanib | VEGFR1, VEGFR2, VEGFR3, CSF1R, FLT3, KIT | RTK signaling | 41 | 210 | -9.32 | 98.68 | 0.490 | 0.941 | 0.987 | **0.701** | 0.157 | 5331 |
| 127 | GSK269962A | ROCK1, ROCK2 | Cytoskeleton | 75 | 71 | -3.09 | 14.29 | 0.576 | 0.869 | 0.983 | **0.701** | 0.105 | 777 |
| 184 | BMS-754807 | IGF1R, IR | IGFR signaling | 196 | 213 | -1.46 | 4.87 | 0.455 | 0.711 | 0.932 | **0.699** | 0.087 | 2480 |
| 62 | BMS-536924 | IGF1R, IR | IGFR signaling | 109 | 68 | -1.28 | 3.81 | 0.507 | 0.768 | 0.972 | **0.699** | 0.055 | 770 |
| 1021 | Axitinib | PDGFR, KIT, VEGFR | RTK signaling | 64 | 178 | -4.85 | 36.32 | 0.483 | 0.841 | 0.979 | **0.699** | 0.045 | 2174 |
| 302 | PI-103 | PI3Kalpha, DAPK3, CLK4, PIM3, HIPK2 | PI3K/MTOR signaling | 470 | 200 | -0.49 | 2.07 | 0.444 | 0.703 | 0.927 | **0.698** | 0.044 | 4909 |
| 1264 | SGC0946 | DOT1L | Chromatin histone methylation | 62 | 221 | -6.21 | 70.14 | 0.268 | 0.965 | 0.987 | **0.696** | 0.023 | 3794 |
| 1036 | PLX-4720 | BRAF | ERK MAPK signaling | 78 | 137 | -4.01 | 22.64 | 0.552 | 0.855 | 0.985 | **0.696** | 0.079 | 2852 |
| 166 | FTI-277 | Farnesyl-transferase (FNTA) | Other | 78 | 163 | -4.36 | 33.25 | 0.413 | 0.937 | 0.987 | **0.695** | 0.025 | 5060 |
| 1142 | HG-5-113-01 | LOK, LTK, TRCB, ABL(T315I) | Other | 45 | 128 | -0.66 | 4.33 | 0.249 | 0.642 | 0.845 | **0.695** | 0.120 | 1727 |
| 178 | BAY-61-3606 | SYK | Other, kinases | 109 | 206 | -1.13 | 4.58 | 0.425 | 0.685 | 0.950 | **0.694** | 0.032 | 3027 |
| 63 | BMS-509744 | ITK | Other | 78 | 85 | -1.28 | 4.52 | 0.405 | 0.846 | 0.965 | **0.689** | 0.068 | 589 |
| 1236 | UNC0638 | G9a and GLP methyltransferases | Chromatin histone methylation | 77 | 304 | -0.95 | 7.86 | 0.367 | 0.727 | 0.909 | **0.688** | 0.083 | 3127 |
| 150 | Bicalutamide | AR | Hormone-related | 40 | 214 | -6.74 | 70.62 | 0.249 | 0.970 | 0.988 | **0.687** | 0.058 | 3348 |
| 199 | Pazopanib | CSF1R, KIT, PDGFRA, PDGFRB | RTK signaling | 115 | 139 | -3.28 | 19.36 | 0.526 | 0.838 | 0.984 | **0.687** | 0.040 | 1688 |
| 5 | Sunitinib | PDGFR, KIT, VEGFR, FLT3, RET, CSF1R | RTK signaling | 46 | 97 | -2.86 | 14.41 | 0.480 | 0.744 | 0.964 | **0.686** | 0.064 | 2140 |
| 1028 | VX-702 | p38 | JNK and p38 signaling | 58 | 143 | -6.98 | 77.05 | 0.419 | 0.938 | 0.988 | **0.686** | 0.028 | 1336 |
| 175 | PAC-1 | Procaspase-3, Procaspase-7 | Apoptosis regulation | 125 | 125 | -1.33 | 4.64 | 0.475 | 0.847 | 0.984 | **0.684** | 0.030 | 3499 |
| 1219 | PFI-1 | BRD4 | Chromatin other | 140 | 217 | -0.92 | 3.92 | 0.347 | 0.790 | 0.938 | **0.682** | 0.031 | 3346 |
| 299 | OSI-027 | MTORC1, MTORC2 | PI3K/MTOR signaling | 471 | 192 | -0.28 | 1.80 | 0.396 | 0.632 | 0.875 | **0.682** | 0.026 | 4072 |
| 1025 | SB216763 | GSK3A, GSK3B | WNT signaling | 157 | 118 | -2.39 | 9.73 | 0.428 | 0.957 | 0.989 | **0.681** | 0.059 | 3329 |
| 1066 | AZD6482 | PI3Kbeta | PI3K/MTOR signaling | 163 | 180 | -1.81 | 6.88 | 0.480 | 0.821 | 0.974 | **0.680** | 0.047 | 3979 |
| 255 | CP724714 | ERBB2 | EGFR signaling | 62 | 187 | -6.63 | 62.59 | 0.458 | 0.915 | 0.985 | **0.679** | 0.078 | 1578 |
| 1005 | Cisplatin | DNA crosslinker | DNA replication | 87 | 195 | -1.63 | 7.57 | 0.396 | 0.788 | 0.955 | **0.679** | 0.083 | 2479 |
| 110 | Seliciclib | CDK2, CDK7, CDK9 | Cell cycle | 71 | 96 | -1.13 | 4.27 | 0.379 | 0.900 | 0.975 | **0.679** | 0.099 | 452 |
| 298 | OSI-930 | KIT | RTK signaling | 61 | 171 | -7.24 | 88.08 | 0.436 | 0.887 | 0.982 | **0.678** | 0.065 | 4231 |
| 153 | Midostaurin | PKC, PPK, FLT1, c-FGR, others | Other | 128 | 202 | -1.59 | 6.69 | 0.429 | 0.804 | 0.964 | **0.676** | 0.020 | 2965 |
| 83 | JW-7-52-1 | MTOR | PI3K/MTOR signaling | 129 | 106 | -0.14 | 1.98 | 0.341 | 0.498 | 0.808 | **0.676** | 0.041 | 1052 |
| 156 | AZD6482 | PI3Kbeta | PI3K/MTOR signaling | 118 | 167 | -2.45 | 11.35 | 0.496 | 0.795 | 0.977 | **0.675** | 0.029 | 2901 |
| 1218 | JQ1 | BRD2, BRD3, BRD4, BRDT | Chromatin other | 154 | 154 | -3.19 | 14.20 | 0.554 | 0.940 | 0.986 | **0.674** | 0.036 | 4600 |
| 37 | Crizotinib | MET, ALK, ROS1 | RTK signaling | 47 | 72 | -4.55 | 27.97 | 0.531 | 0.920 | 0.987 | **0.671** | 0.121 | 896 |
| 229 | Enzastaurin | PKCB | Other, kinases | 285 | 192 | -1.36 | 4.22 | 0.502 | 0.811 | 0.970 | **0.671** | 0.033 | 1781 |
| 140 | Vinorelbine | Microtubule destabiliser | Mitosis | 121 | 221 | -0.21 | 2.03 | 0.353 | 0.501 | 0.876 | **0.670** | 0.032 | 1152 |
| 35 | NVP-TAE684 | ALK | RTK signaling | 54 | 89 | -2.31 | 10.63 | 0.475 | 0.696 | 0.956 | **0.668** | 0.100 | 1077 |
| 1067 | CCT007093 | PPM1D | Other | 86 | 162 | -3.06 | 16.30 | 0.388 | 0.944 | 0.988 | **0.667** | 0.039 | 3637 |
| 201 | Epothilone B | Microtubule stabiliser | Mitosis | 135 | 213 | -0.30 | 2.15 | 0.364 | 0.409 | 0.853 | **0.667** | 0.027 | 1369 |
| 1143 | HG-5-88-01 | EGFR, ADCK4 | Other, kinases | 60 | 90 | -3.47 | 23.77 | 0.472 | 0.826 | 0.976 | **0.667** | 0.073 | 1810 |
| 1016 | Temsirolimus | MTOR | PI3K/MTOR signaling | 209 | 174 | -1.25 | 4.31 | 0.438 | 0.696 | 0.933 | **0.666** | 0.040 | 5381 |
| 328 | SNX-2112 | HSP90 | Protein stability and degradation | 609 | 226 | -0.22 | 1.75 | 0.396 | 0.793 | 0.875 | **0.666** | 0.047 | 5611 |
| 249 | Cabozantinib | VEGFR, MET, RET, KIT, FLT1, FLT3, FLT4, TIE2,AXL | Other, kinases | 67 | 202 | -4.32 | 29.60 | 0.490 | 0.812 | 0.977 | **0.666** | 0.102 | 2048 |
| 87 | GW843682X | PLK1 | Cell cycle | 40 | 116 | -0.10 | 1.95 | 0.364 | 0.435 | 0.854 | **0.664** | 0.097 | 1533 |
| 32 | Tozasertib | AURKA, AURKB, AURKC, others | Mitosis | 112 | 55 | -1.50 | 4.36 | 0.576 | 0.794 | 0.984 | **0.662** | 0.098 | 1057 |
| 177 | GSK650394 | SGK2, SGK3 | Other | 190 | 169 | -1.44 | 5.02 | 0.503 | 0.780 | 0.979 | **0.662** | 0.035 | 236 |
| 1192 | GSK269962A | ROCK1, ROCK2 | Cytoskeleton | 154 | 198 | -2.19 | 8.84 | 0.472 | 0.787 | 0.961 | **0.661** | 0.024 | 1431 |
| 1091 | BMS-536924 | IGF1R, IR | Unclassified | 212 | 203 | -1.52 | 5.32 | 0.446 | 0.864 | 0.969 | **0.660** | 0.052 | 2455 |
| 1054 | Palbociclib | CDK4, CDK6 | Cell cycle | 137 | 165 | -1.32 | 4.72 | 0.492 | 0.735 | 0.977 | **0.659** | 0.063 | 4291 |
| 207 | AS601245 | JNK1, JNK2, JNK2 | JNK and p38 signaling | 96 | 221 | -0.95 | 4.39 | 0.345 | 0.754 | 0.928 | **0.656** | 0.076 | 1629 |
| 1053 | MK-2206 | AKT1, AKT2 | PI3K/MTOR signaling | 144 | 179 | -1.44 | 5.22 | 0.455 | 0.752 | 0.955 | **0.654** | 0.023 | 2613 |
| 197 | Bryostatin 1 | PKC | Other, kinases | 125 | 145 | -3.37 | 18.81 | 0.454 | 0.939 | 0.988 | **0.653** | 0.047 | 2774 |
| 158 | PF-562271 | FAK, FAK2 | Cytoskeleton | 74 | 212 | -1.82 | 9.37 | 0.388 | 0.838 | 0.967 | **0.653** | 0.067 | 1351 |
| 41 | S-Trityl-L-cysteine | KIF11 | Mitosis | 55 | 102 | -0.16 | 1.93 | 0.363 | 0.634 | 0.908 | **0.652** | 0.093 | 1611 |
| 9 | MG-132 | Proteasome, CAPN1 | Protein stability and degradation | 41 | 101 | -0.76 | 3.40 | 0.403 | 0.661 | 0.937 | **0.652** | 0.130 | 1410 |
| 1058 | Pictilisib | PI3K (class 1) | PI3K/MTOR signaling | 133 | 184 | -1.18 | 4.40 | 0.427 | 0.662 | 0.938 | **0.652** | 0.042 | 2738 |
| 1057 | Dactolisib | PI3K (Class 1), MTORC1, MTORC2 | PI3K/MTOR signaling | 86 | 209 | -0.64 | 3.74 | 0.297 | 0.538 | 0.835 | **0.651** | 0.057 | 2706 |
| 1529 | Pevonedistat | NAE | Other | 189 | 148 | -1.04 | 3.70 | 0.439 | 0.785 | 0.957 | **0.649** | 0.027 | 1893 |
| 1530 | PFI-3 | SMARCA2, SMARCA4, PB1 | Other | 45 | 176 | -5.35 | 49.19 | 0.311 | 0.951 | 0.986 | **0.646** | 0.104 | 1518 |
| 34 | Imatinib | ABL, KIT, PDGFR | RTK signaling | 22 | 76 | -5.67 | 37.81 | 0.593 | 0.922 | 0.990 | **0.646** | 0.090 | 3418 |
| 1043 | JNK Inhibitor VIII | JNK | JNK and p38 signaling | 101 | 162 | -3.17 | 17.57 | 0.392 | 0.951 | 0.988 | **0.645** | 0.043 | 5541 |
| 1230 | IOX2 | EGLN1 | Other | 50 | 193 | -10.55 | 188.01 | 0.387 | 0.939 | 0.987 | **0.645** | 0.069 | 5438 |
| 1170 | CCT-018159 | HSP90 | Protein stability and degradation | 42 | 199 | -1.56 | 11.95 | 0.352 | 0.802 | 0.959 | **0.642** | 0.086 | 2131 |
| 54 | CGP-082996 | CDK4 | Cell cycle | 34 | 80 | -1.79 | 8.71 | 0.438 | 0.847 | 0.981 | **0.642** | 0.131 | 1130 |
| 1038 | NU7441 | DNAPK | Genome integrity | 123 | 152 | -1.75 | 7.13 | 0.405 | 0.918 | 0.984 | **0.640** | 0.062 | 4260 |
| 1261 | rTRAIL | TRAIL receptor agonist | Apoptosis regulation | 199 | 126 | -2.93 | 12.98 | 0.590 | 0.911 | 0.986 | **0.638** | 0.063 | 1507 |
| 11 | Paclitaxel | Microtubule stabiliser | Mitosis | 71 | 94 | -0.37 | 2.07 | 0.412 | 0.521 | 0.922 | **0.636** | 0.065 | 455 |
| 1052 | RO-3306 | CDK1 | Cell cycle | 142 | 129 | -2.24 | 9.12 | 0.472 | 0.919 | 0.987 | **0.634** | 0.039 | 4423 |
| 147 | NSC-87877 | SHP-1 (PTPN6), SHP-2 (PTPN11) | Other | 91 | 181 | -4.08 | 26.36 | 0.430 | 0.942 | 0.987 | **0.634** | 0.046 | 2158 |
| 293 | Amuvatinib | KIT, PDGFRA, FLT3 | Other, kinases | 184 | 191 | -2.65 | 10.39 | 0.529 | 0.834 | 0.967 | **0.632** | 0.037 | 1559 |
| 1069 | EHT-1864 | RAC1, RAC2, RAC3 | Cytoskeleton | 99 | 187 | -1.62 | 6.58 | 0.379 | 0.898 | 0.979 | **0.631** | 0.050 | 672 |
| 1239 | YK-4-279 | RNA helicase A | Other | 100 | 217 | -0.06 | 2.49 | 0.287 | 0.622 | 0.839 | **0.626** | 0.042 | 2707 |
| 71 | Pyrimethamine | Dihydrofolate reductase (DHFR) | Other | 96 | 71 | -1.57 | 5.22 | 0.511 | 0.832 | 0.980 | **0.624** | 0.084 | 599 |
| 186 | Bexarotene | Retinioic X receptor (RXR) agonist | Other | 50 | 167 | -4.98 | 51.83 | 0.464 | 0.847 | 0.983 | **0.623** | 0.111 | 2449 |
| 164 | JQ12 | HDAC1, HDAC2 | Chromatin histone acetylation | 176 | 183 | -0.91 | 3.45 | 0.422 | 0.645 | 0.938 | **0.618** | 0.043 | 346 |
| 3 | Rapamycin | MTORC1 | PI3K/MTOR signaling | 80 | 58 | -1.78 | 5.29 | 0.578 | 0.736 | 0.978 | **0.615** | 0.047 | 995 |
| 17 | Cyclopamine | SMO | Other | 92 | 58 | -2.27 | 8.70 | 0.466 | 0.953 | 0.988 | **0.615** | 0.039 | 986 |
| 1375 | Temozolomide | DNA alkylating agent | DNA replication | 75 | 178 | -5.64 | 54.51 | 0.378 | 0.943 | 0.986 | **0.613** | 0.053 | 1962 |
| 291 | KIN001-266 | MAP3K8 | Other, kinases | 109 | 297 | -0.89 | 4.93 | 0.325 | 0.807 | 0.914 | **0.612** | 0.089 | 877 |
| 111 | Salubrinal | EIF2A | Other | 46 | 93 | -0.99 | 3.99 | 0.383 | 0.837 | 0.966 | **0.610** | 0.097 | 1127 |
| 173 | FH535 | PPARgamma, PPARdelta | WNT signaling | 99 | 234 | -0.29 | 3.02 | 0.265 | 0.492 | 0.782 | **0.609** | 0.051 | 3728 |
| 185 | Linsitinib | IGF1R | IGFR signaling | 164 | 131 | -2.40 | 9.25 | 0.581 | 0.856 | 0.986 | **0.604** | 0.038 | 3050 |
| 172 | Embelin | XIAP | Apoptosis regulation | 83 | 233 | -0.22 | 2.75 | 0.276 | 0.659 | 0.866 | **0.603** | 0.050 | 2122 |
| 1042 | Doramapimod | p38, JNK2 | JNK and p38 signaling | 76 | 130 | -3.26 | 20.95 | 0.458 | 0.910 | 0.988 | **0.603** | 0.038 | 4492 |
| 60 | BI-2536 | PLK1, PLK2, PLK3 | Cell cycle | 60 | 110 | -0.35 | 2.18 | 0.379 | 0.593 | 0.899 | **0.600** | 0.125 | 956 |
| 133 | Doxorubicin | Anthracycline | DNA replication | 113 | 230 | -0.19 | 2.21 | 0.314 | 0.372 | 0.788 | **0.599** | 0.041 | 689 |
| 190 | Bleomycin | dsDNA break induction | DNA replication | 618 | 257 | -0.04 | 1.75 | 0.353 | 0.730 | 0.773 | **0.599** | 0.029 | 1455 |
| 269 | NSC-207895 | MDM4 | p53 pathway | 159 | 173 | -1.83 | 7.24 | 0.503 | 0.824 | 0.982 | **0.599** | 0.030 | 1390 |
| 1194 | SB505124 | ALK4, ALK5 | RTK signaling | 108 | 167 | -3.41 | 20.95 | 0.419 | 0.929 | 0.985 | **0.598** | 0.067 | 996 |
| 1049 | PD173074 | FGFR1, FGFR3 | RTK signaling | 50 | 126 | -5.27 | 41.93 | 0.528 | 0.876 | 0.988 | **0.598** | 0.058 | 1336 |
| 1029 | Motesanib | VEGFR, RET, KIT, PDGFR | RTK signaling | 33 | 146 | -9.21 | 140.88 | 0.478 | 0.893 | 0.988 | **0.596** | 0.118 | 5365 |
| 1017 | Olaparib | PARP1, PARP2 | Genome integrity | 81 | 118 | -3.48 | 22.68 | 0.458 | 0.912 | 0.987 | **0.595** | 0.052 | 1900 |
| 1133 | Serdemetan | MDM2 | p53 pathway | 84 | 231 | -1.63 | 8.80 | 0.341 | 0.797 | 0.941 | **0.594** | 0.081 | 2918 |
| 1502 | Bicalutamide | AR | Hormone-related | 31 | 166 | -8.14 | 102.99 | 0.352 | 0.933 | 0.986 | **0.593** | 0.028 | 2772 |
| 1018 | Veliparib | PARP1, PARP2 | Genome integrity | 81 | 166 | -4.42 | 33.68 | 0.354 | 0.957 | 0.988 | **0.587** | 0.112 | 3596 |
| 167 | OSU-03012 | PDK1 (PDPK1) | Other, kinases | 119 | 231 | -0.52 | 3.15 | 0.292 | 0.608 | 0.852 | **0.581** | 0.021 | 2378 |
| 1031 | Elesclomol | HSP90 | Protein stability and degradation | 96 | 247 | -0.14 | 2.17 | 0.355 | 0.401 | 0.817 | **0.579** | 0.065 | 3162 |
| 205 | Avagacestat | Amyloid beta20, Amyloid beta40 | Other | 107 | 196 | -3.48 | 20.09 | 0.291 | 0.970 | 0.988 | **0.576** | 0.133 | 2491 |
| 202 | GSK1904529A | IGF1R, IR | IGFR signaling | 97 | 160 | -3.37 | 20.96 | 0.381 | 0.950 | 0.988 | **0.575** | 0.054 | 4268 |
| 1046 | Wee1 Inhibitor | WEE1, CHEK1 | Cell cycle | 41 | 143 | -3.06 | 21.77 | 0.458 | 0.847 | 0.984 | **0.572** | 0.042 | 1691 |
| 89 | Parthenolide | HDAC1 | Chromatin histone acetylation | 49 | 60 | -1.61 | 6.06 | 0.494 | 0.862 | 0.987 | **0.571** | 0.109 | 770 |
| 163 | JQ1 | BRD2, BRD3, BRD4, BRDT | Chromatin other | 99 | 204 | -0.52 | 2.57 | 0.368 | 0.544 | 0.899 | **0.571** | 0.035 | 1653 |
| 1020 | Lenalidomide | CRBN | Protein stability and degradation | 57 | 158 | -6.36 | 65.56 | 0.395 | 0.944 | 0.988 | **0.568** | 0.121 | 3312 |
| 151 | QS11 | ARFGAP1 | Other | 158 | 181 | -1.55 | 5.72 | 0.478 | 0.835 | 0.979 | **0.568** | 0.059 | 829 |
| 1023 | GW441756 | NTRK1 | RTK signaling | 80 | 137 | -4.68 | 36.55 | 0.532 | 0.885 | 0.987 | **0.568** | 0.032 | 4679 |
| 193 | GW-2580 | CSF1R | RTK signaling | 36 | 205 | -15.05 | 319.43 | 0.357 | 0.943 | 0.987 | **0.564** | 0.129 | 3037 |
| 1164 | XMD8-92 | MAPK7 | Other, kinases | 54 | 119 | -0.76 | 3.39 | 0.342 | 0.842 | 0.956 | **0.560** | 0.093 | 1282 |
| 1072 | Avagacestat | Amyloid beta20, Amyloid beta40 | Other | 67 | 139 | -4.24 | 34.27 | 0.458 | 0.911 | 0.988 | **0.556** | 0.111 | 4456 |
| 6 | PHA-665752 | MET | RTK signaling | 26 | 75 | -7.43 | 73.26 | 0.495 | 0.927 | 0.989 | **0.550** | 0.175 | 1592 |
| 1262 | UNC1215 | L3MBTL3 | Chromatin other | 84 | 198 | -3.25 | 21.54 | 0.235 | 0.970 | 0.987 | **0.550** | 0.062 | 2706 |
| 1199 | Tamoxifen | ESR1 | Hormone-related | 46 | 169 | -5.82 | 60.34 | 0.380 | 0.926 | 0.986 | **0.546** | 0.171 | 1217 |
| 192 | LFM-A13 | BTK | Other, kinases | 118 | 172 | -2.26 | 11.24 | 0.376 | 0.939 | 0.985 | **0.541** | 0.038 | 2117 |
| 1039 | SL0101 | RSK, AURKB, PIM1, PIM3 | Other | 111 | 163 | -3.30 | 20.36 | 0.359 | 0.957 | 0.987 | **0.538** | 0.063 | 2802 |
| 86 | A-443654 | AKT1, AKT2, AKT3 | PI3K/MTOR signaling | 43 | 111 | -0.33 | 2.75 | 0.325 | 0.595 | 0.862 | **0.534** | 0.120 | 136 |
| 204 | Tipifarnib | Farnesyl-transferase (FNTA) | Other | 186 | 198 | -0.78 | 3.21 | 0.367 | 0.554 | 0.868 | **0.530** | 0.030 | 1705 |
| 1527 | Pictilisib | PI3K (class 1) | PI3K/MTOR signaling | 114 | 207 | -0.84 | 3.63 | 0.361 | 0.635 | 0.902 | **0.526** | 0.021 | 2963 |
| 1158 | XMD11-85h | BRSK2, FLT4, MARK4, PRKCD, RET, SPRK1 | Other | 85 | 87 | -1.63 | 6.48 | 0.381 | 0.919 | 0.982 | **0.516** | 0.064 | 1903 |
| 91 | GSK319347A | IKK | Other | 45 | 84 | -4.09 | 29.39 | 0.345 | 0.965 | 0.989 | **0.512** | 0.067 | 1688 |
| 1129 | PF-4708671 | S6K1 | PI3K/MTOR signaling | 73 | 179 | -3.08 | 19.25 | 0.408 | 0.896 | 0.981 | **0.506** | 0.087 | 207 |
| 1033 | Vismodegib | SMO | Other | 117 | 147 | -2.86 | 14.32 | 0.384 | 0.950 | 0.987 | **0.505** | 0.037 | 1213 |
| 341 | Selisistat | SIRT1 | Chromatin histone acetylation | 63 | 197 | -5.51 | 49.44 | 0.276 | 0.965 | 0.987 | **0.436** | 0.082 | 2289 |
| 266 | Zibotentan | Endothelin-1 receptor (EDNRA) | Other | 50 | 223 | -8.37 | 131.85 | 0.261 | 0.969 | 0.988 | **0.422** | 0.100 | 2574 |
